# Andes virus from the *MV Hondius* outbreak is highly pathogenic in hamsters and airborne shed

**DOI:** 10.64898/2026.10.12.758426

**Authors:** Melanie Rissmann, Maarten F. Wilbrink, Vanessa Herder, Kim Handrejk, Fabien Filaire, Georgios Ilia, Wai Wai Jones, Luca M. Zaeck, Debby van Eck Schipper, Vincent P. Vaes, Sandhya Shrestha, Johan Reimerink, Laura L.A. van Dijk, Mark Power, Merel R. te Marvelde, Dennis de Meulder, Hayley Cassidy, Koen Wijnans, Sharona Sacharoen–Rugebregt, Alba Grifoni, Alessandro Sette, Bas B. Oude Munink, Chantal B.E.M. Reusken, Bart L. Haagmans, Rory D. de Vries, Sander Herfst, Massimo Palmarini

## Abstract

In May 2026, an outbreak of hantavirus pulmonary syndrome (HPS) caused by Andes virus (ANDV) was detected aboard the cruise ship *MV Hondius,* sparking public health measures in multiple countries. ANDV infections in humans are mostly sporadic, most often linked to exposure to infected rodents in South America, although small clusters of human cases, like the 2026 outbreak on the *MV Hondius* occur as well. It is not known whether virological factors may contribute to this distinct epidemiology, and no phenotypic characterisation of the newly emerged ANDV (ANDV_2026_) has been carried out to date. Here, using an ANDV isolate from a presymptomatic patient from the *MV Hondius*, we studied pathogenesis, identified sites of viral replication, characterized innate and adaptive immune responses, and quantified shedding in golden Syrian hamsters. ANDV_2026_ proved to be highly pathogenic in hamsters, causing severe pulmonary disease reminiscent of HPS. ANDV_2026_ triggered a delayed type-I IFN response and inflammatory response in the lungs consistent with the key pathological features of HPS, and we detected viral replication in both respiratory and extra-respiratory tissues, including the brain. Infected animals mounted neutralising antibodies and T cell responses, the latter directed predominantly against the L and N proteins. Importantly, ANDV_2026_-infected animals shed virus through multiple routes, and we detected infectious virus in air samplers connected to cages housing ANDV_2026_-infected hamsters. In contrast, control hamsters infected with a rodent-derived ANDV isolate (ANDV_9717869_) shed little virus and air samples were consistently negative. These findings consolidate established features of ANDV pathogenesis, while revealing properties not previously described. Phenotypic characterisation of patient-derived ANDV isolates in animal models can strengthen outbreak preparedness, identify markers of disease severity, and support the development of medical countermeasures.

## INTRODUCTION

On the 2^nd^ of May 2026, a cluster of severe respiratory disease aboard the Dutch cruise ship *MV Hondius* was notified to the World Health organization (WHO)^1–3^. Soon after, ANDV was diagnosed as the cause of the outbreak, which overall led to 13 reported cases, including three deaths^4^. Confirmed cases were all aboard the *MV Hondius*, although the outbreak led to monitoring of 653 individuals in 33 countries around the globe.

ANDV is classified within the family *Orthohantaviridae*^5–8^. These are viruses that cause two distinct disease complexes: the hemorrhagic fever with renal syndrome (HFRS) in Europe and Asia, and the hantavirus pulmonary syndrome (HPS) in the Americas, caused among others by Sin Nombre virus (SNV) and Andes virus (ANDV), respectively^9^. HPS is usually presents in four distinct clinical phases: febrile, cardiopulmonary, diuretic and convalescent^10,11^. Most cases are characterized by pulmonary edema driven by increased vascular permeability, terminally leading to cardiogenic shock and death^12^. The case fatality rate of the *MV Hondius* outbreak was 23%, lower than for previously reported ANDV clusters. Direct comparison is difficult however, due to the relatively low number of cases in this outbreak and early access to medical care for many of the individuals affected^13–15^.

ANDV is endemic in Southern Argentina and Chile, where it causes seasonal outbreaks every year^9,16^. Humans are believed to become infected mostly after exposure to aerosolized infectious excretions of the rodent reservoir, the long-tailed rice rat (*Oligoryzomys longicaudatus*)^5,6,17^. Andes virus (ANDV) was first detected in 1995, when it caused an outbreak of 16 cases and nine deaths^18^. This outbreak and subsequent ones suggested that ANDV can be capable of human-to-human transmission, with the main risk factor being in close contact with infected individuals in enclosed settings^18,19^. Some cases have also been described without any documented direct contact, hinting at a respiratory transmission route, though available data cannot distinguish airborne shedding of infectious virus from contamination of the surrounding environment^20^. On the *MV Hondius,* patients were in direct contact with each other, suggesting ANDV human-to-human transmission. One potential asymptomatic case of ANDV infection in a health-care worker with low-level viraemia was also reported, despite only indirect exposure^21–23^.

Few studies have phenotyped ANDV isolates obtained directly from human patients^21–23^, and fewer still have done so *in vivo* to define key viral characteristics such as virulence and host responses^22,24^. This reflects the sporadic nature of HPS and its restricted geographical distribution. ANDV is also notoriously difficult to isolate, as infection is typically diagnosed once patients already developed neutralising antibodies. As a result, our understanding of the biological differences between ANDV isolates associated with different cases and outbreaks is incomplete.

Here, we characterised the ANDV isolate associated with the *MV Hondius* outbreak (ANDV/2026/hondius-NL-12, henceforth ANDV_2026_). Using the golden Syrian hamster model, we studied viral pathogenesis, replication sites, and the host innate and adaptive immune responses. We also analysed virus shedding from infected animals and sampled air from the cages of infected hamsters for ANDV RNA and infectious virus. Throughout our study, the human isolate ANDV_2026_ was compared with a rodent-derived ANDV strain (ANDV_9717869_). Altogether, in vivo phenotyping contributes to the continuous risk assessment of emerging viruses of global health concern.

## RESULTS

### ANDV_2026_ causes severe, systemic infection in hamsters

Golden Syrian hamsters were intranasally inoculated with a low dose (200 TCID_50_/hamster) of either ANDV_2026_ (n=18) or ANDV_9717869_ (n=18). ANDV_2026_ was isolated from whole-blood sample collected from a pre-symptomatic patient during the 2026 ANDV outbreak linked to the cruise ship *MV Hondius*^3^. Backtitration of the inocula revealed that the animals received a dose of 28 TCID50 of ANDV_2026_ and 89 TCID50 of ANDV_9717869_. Six animals per group were sacrificed at 3, 7 and 14 days post-inoculation (dpi), or earlier when reaching humane endpoints. Sixty percent of the hamsters inoculated with ANDV_2026_ reached the humane endpoint between 12 and 14 dpi (**Fig 1a**). Symptoms included lethargy, ruffled fur, decreased mobility, and severe respiratory distress. In general, animals reached humane endpoints within 24-72 hours after the onset of symptoms. The development of clinical symptoms was also accompanied by a loss of body weight of up to 9% in three days (**Fig. 1b**). At the terminal stage of the experiment (12-14 dpi), the lung-to-bodyweight ratio of ANDV_2026_ inoculated animals was significantly increased when compared to both mock- and ANDV_9717869_-inoculated animals (**Fig. 1c**). These values reflected the macroscopic lung lesions observed in ANDV_2026_ inoculated animals, characterized by increased size, rounded edges and darkened colour with severe, diffuse lung oedema (**Fig. 1d**), as described previously for severe ANDV infections in hamsters^25–27^.

**Figure 1.**
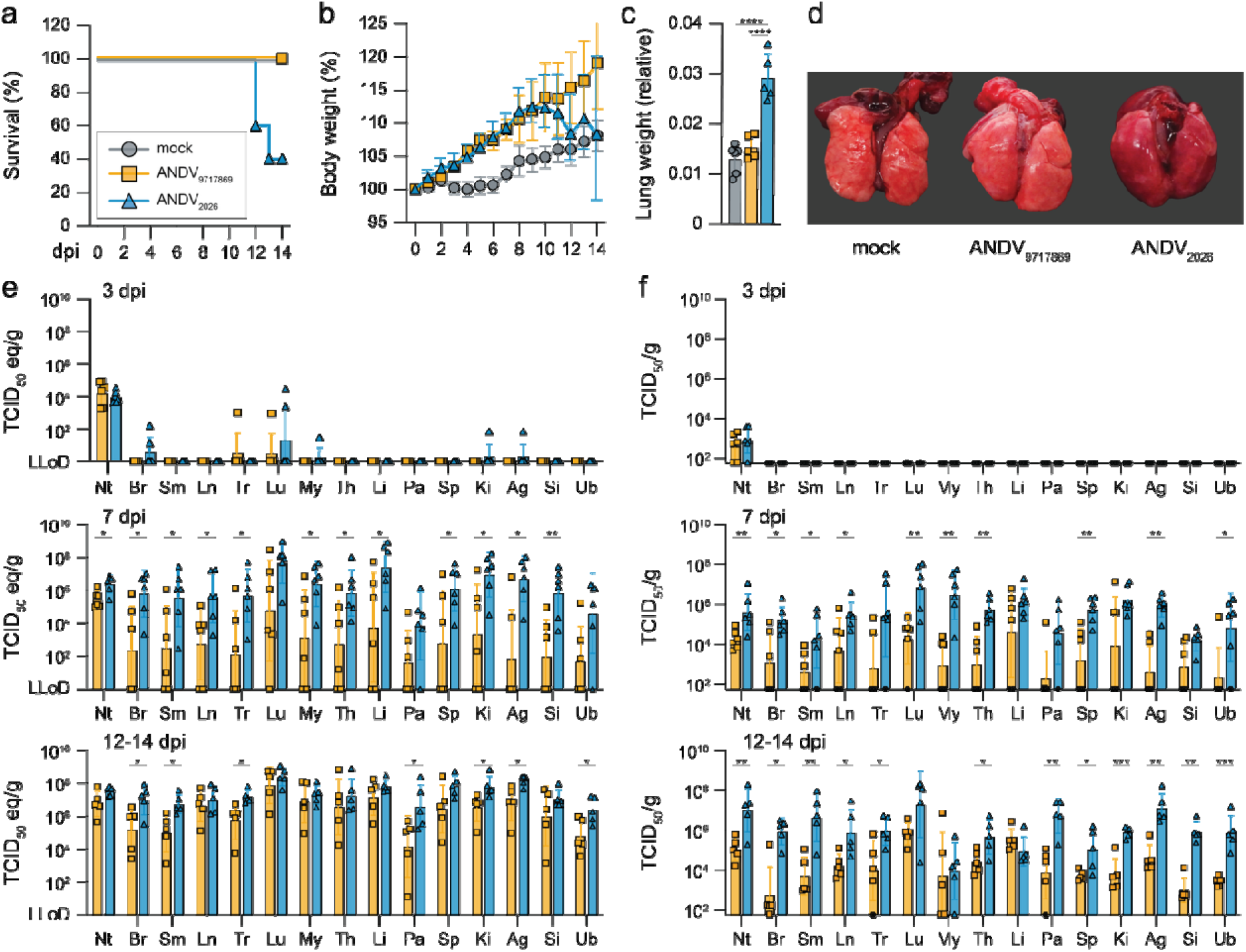
Severe disease and systemic spread of ANDV_2026_ in experimentally inoculated hamsters. (**a**) Kaplan-Meier curve of hamsters experimentally inoculated with ANDV. (**b**) Body weight relative to starting weight at 0 dpi. Data shown as mean line graph ± SD. (**c**) Lung-to-body weight ratios at 12-14 dpi. Data shown as mean ± SD; symbols represent individual hamsters. Data was compared by one-way ANOVA with multiple comparisons; **** p<0.0001. (**d**) Representative macroscopic images of lungs from experimentally inoculated hamsters taken at 14 dpi. (**e**) Quantification of viral genome in tissues measured by qRT-PCR on 3, 7 and 12-14 dpi. Data shown as geometric mean ± geometric SD; symbols represent individual hamsters. Log-transformed data was compared by unpaired t test or Mann-Whitney test for normal or non-normal distributed data, respectively; * p<0.05, ** p<0.01. (**f**) Quantification of viral load tested in tissues measured by TCID_50_ titration on VeroE6 cells on 3, 7 and 12-14 dpi. Only qRT-PCR positive samples were subjected to virus titrations, not tested samples are indicated by a crossed diamond. Data shown as geometric mean ± geometric SD; symbols represent individual hamsters. Log-transformed data was compared by unpaired t test or Mann-Whitney test for normal or non-normal distributed data, respectively; * p<0.05, ** p<0.01, *** p<0.001. Abbreviations: nasal turbinate (Nt); brain (Br); submandibular gland (Sm); cervical lymph node (Ln); trachea (Tr); lung (Lu); myocardium (My); thymus (Th); liver (Li); pancreas (Pa); spleen (Sp); kidney (Ki); adrenal gland (Ag); small intestine (Si); urinary bladder (Ub); lower limit of detection (LLOD); days post inoculation (dpi).

Next, we tested 15 respiratory and non-respiratory tissues for presence of both viral RNA by qRT-PCR and infectious virus by titration (**Fig. 1e,f**). At the early timepoints (3 dpi), viral RNA and infectious virus was only detected in the nasal turbinates with titers essentially identical in both ANDV_2026_- and ANDV_9717869_-inoculated animals. At 7 and 12-14 dpi all tissues contained both viral RNA and infectious virus with titers in most cases higher in ANDV_2026_-infected animals. Interestingly, we observed no significant differences in viral loads in myocardium and liver between ANDV_2026_- and ANDV_9717869_-infected hamsters 12-14 dpi. These tissues were previously reported as the prime targets of virus replication in ANDV_9717869_-infected hamsters^26^. We observed the highest viral titers in the respiratory tract (nasal turbinates and lungs), submandibular salivary glands, pancreas, and adrenal glands collected from ANDV_2026_-infected animals.

### Sites of ANDV replication

Next, we investigated the sites of viral replication using immunohistochemistry (IHC) and RNA *in situ-*hybridi**s**ation (ISH) in formalin-fixed and paraffin embedded samples from inoculated animals. In the lungs, we detected abundant viral RNA (**Fig. 2a**) and protein in ANDV_2026_-infected animals at 7dpi, with widespread distribution in the parenchyma. Quantification of RNA signal showed virus spread in wider areas of the lungs in ANDV_2026_-infected hamsters compared to those infected with ANDV_9717869_ (**Fig. 2b-c**), both at 7 and 14 dpi. Wirus RNA was detectable, both by qualitative assessment of the histological sections and by software-assisted image analysis^28^, in the lung parenchyma and endothelial cells, but not in the bronchial epithelium of all ANDV infected animals at both 7 and 14 dpi (**Fig. 2c-d**). As expected, extrarespiratory organs such as liver, kidney, myocardium, and many others showed abundant viral RNA signal in endothelial cells, while epithelial cells were viral RNA negative (**Fig. 2e**). Importantly, we also detected ANDV_2026_ widely distributed in the brain, including in endothelial cells of blood vessels, meninges and neurons (**Fig. 2e and Fig. S1**).

**Figure 2.**
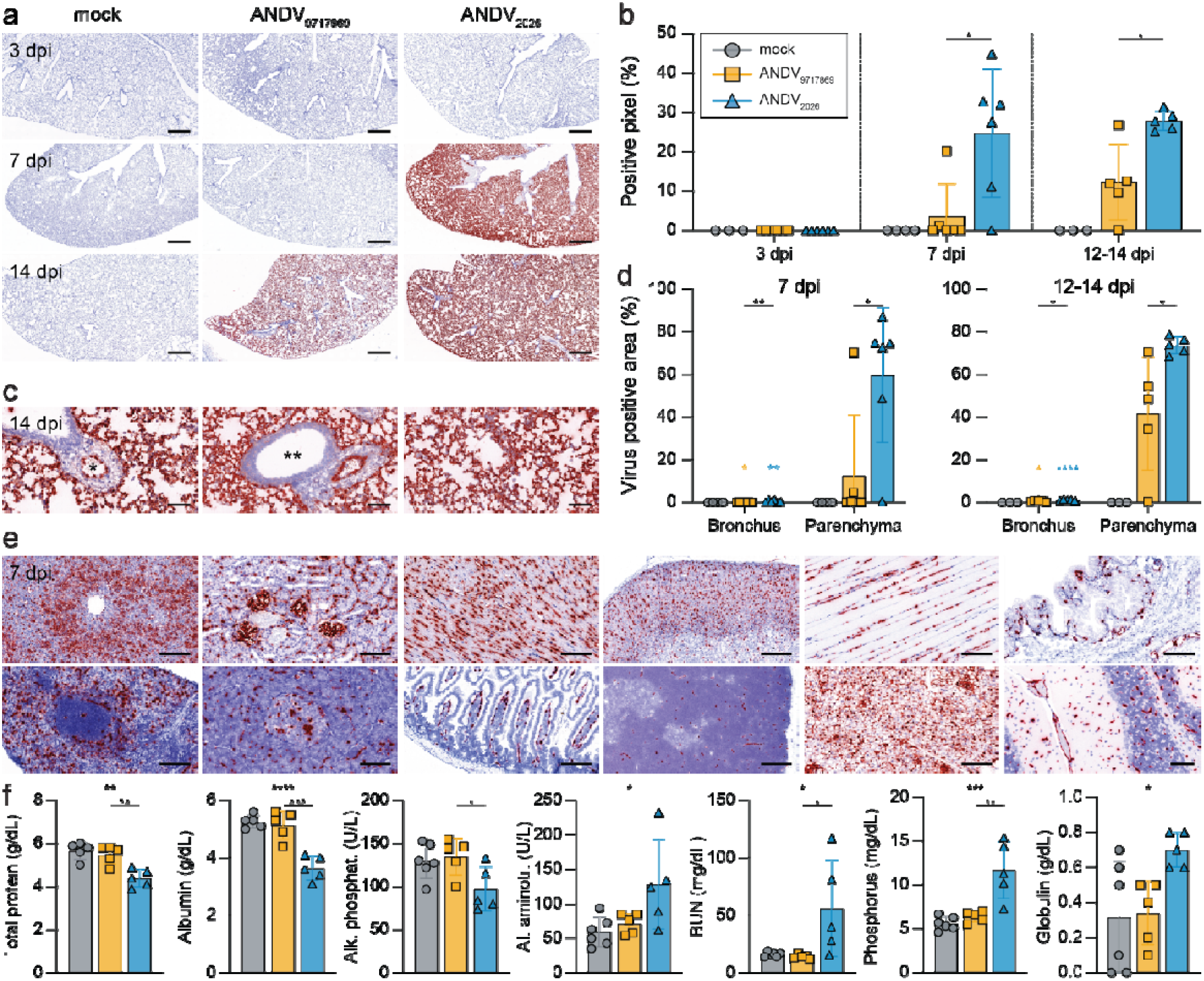
Detection of viral RNA in tissues of ANDV inoculated hamsters. (**a**) *In situ-*hybridisation detecting viral RNA (S-segment) of ANDV_9717869_ and ANDV_2026_ in FFPE lung sections of hamsters experimentally inoculated as indicated. Red signal shows virus positive cells. Bar, 1mm. (**b**) Software-assisted quantification of the virus-positive signal in the whole scanned lung section in all experimentally inoculated animals. Data are expressed as positive pixels and shown as mean ± SD; symbols represent individual hamsters. ANDV_9717869_ and ANDV_2026_ were compared by Mann-Whitney test for non-normal distributed data; * p<0.05. (**c**) Higher magnification of virus-positive areas of a animal inoculated with Andes_2026_. Left panel, * = vessel lumen with strongly positive endothelial cells; middle panel, ** = bronchus lacking viral RNA surrounded by strongly positive lung parenchyma; right panel, area of strongly positiv lung parenchyma. Bar, 100 µm. (**d**) Bar graphs representing software-assisted quantification of the virus positive lun areas separated into bronchus and parenchyma, respectively. Graphs show viral quantification of viral RNA at both 7 and 12-14 dpi. Data shown as mean ± SD; symbols represent individual hamsters. ANDV_9717869_ and ANDV_2026_ were compared by Mann-Whitney test for non-normal distributed data; * p<0.05, ** p<0.01. Colour-coded asterisks denote the difference between bronchus and parenchyma, compared by t test or Wilcoxon rank test for paired normal or non-normal distributed data, respectively. (**e**) Representative micrographs of *in situ* hybridization of viral RNA in FFPE sections of various organs collected from hamsters inoculated with ANDV_2026_ at 7 dpi: top row from left to right: liver (bar, 200 µm), kidney (bar, 100 µm), myocardium (bar, 100 µm), adrenal gland (bar, 100 µm), musculus quadriceps (bar, 100 µm), urinary bladder (bar, 100 µm); bottom row, from left to right: spleen (bar, 100 µm), pancreas (bar, 100 µm), small intestine (bar, 100 µm), thymus (bar, 100 µm), abdominal adipose tissue (bar, 100 µm), and brain (bar, 100 µm). (**f**) Bar graphs showing blood parameters of ANDV inoculated hamsters at 12-14 dpi: total protein, albumin, alkaline phosphatase, alanine-aminotransferase, blood urea nitrogen, phosphorus, globulin (from left to right). Dat shown as mean ± SD; symbols represent individual hamsters. Abbreviations: days post inoculation (dpi); blood ure nitrogen (BUN). Data was compared by one-way ANOVA with multiple comparisons; * p<0.05, ** p<0.01, *** p<0.001, **** p<0.0001.

The widespread infection of endothelial cells throughout all organs was not accompanied by severe histopathological lesions in both groups of animals. Serum chemistry revealed reduced total protein and albumin concentrations in ANDV_2026_-infected hamsters compared to ANDV_9717869_- and mock-infected animals, consistent with endothelial dysfunction and leakage of proteins to the extravascular space (**Fig. 2f**). We found moderately abnormal values for alkaline phosphatase, alanine-aminotransferase, blood urea nitrogen, creatinine and phosphorus in ANDV_2026_-infected animals (in the terminal stages of the disease), consistent with kidney and liver damage, although major histopathological lesions were not observed (**Fig. 2f, Fig. S2**). Serum values of amylase, calcium, glucose, sodium and total bilirubin remained similar between groups (**Fig. S2**).

### Lung pathology in ANDV_2026_-infected hamsters correlates with preferential activation of the type I IFN response

At the latest stages of disease, we detected histopathological lesions throughout the lungs in hamsters infected with ANDV_2026_, similar to severe HPS previously described in this model ^25–27^. Lesions include diffuse alveolar damage, multifocal, lymphoplasmacytic infiltration, alveolar histiocytosis, alveolar oedema, leukocytolastic vasculitis, perivascular oedema, sloughing of bronchial epithelial cells to various degrees and hemorrhages in the vessel (**Fig. 3a-b**). We also detected fibrin accumulation in damaged alveoli in more severe cases (**Fig. 3b**, middle panel). Infiltrates of both B-cells (Pax^+^ cells) and macrophage (Iba-1^+^ cells) infiltrates, but not T-cells (CD3^+^), were observed in the lungs of ANDV_2026_-infected animals (**Fig. 3c-d**).

**Figure 3.**
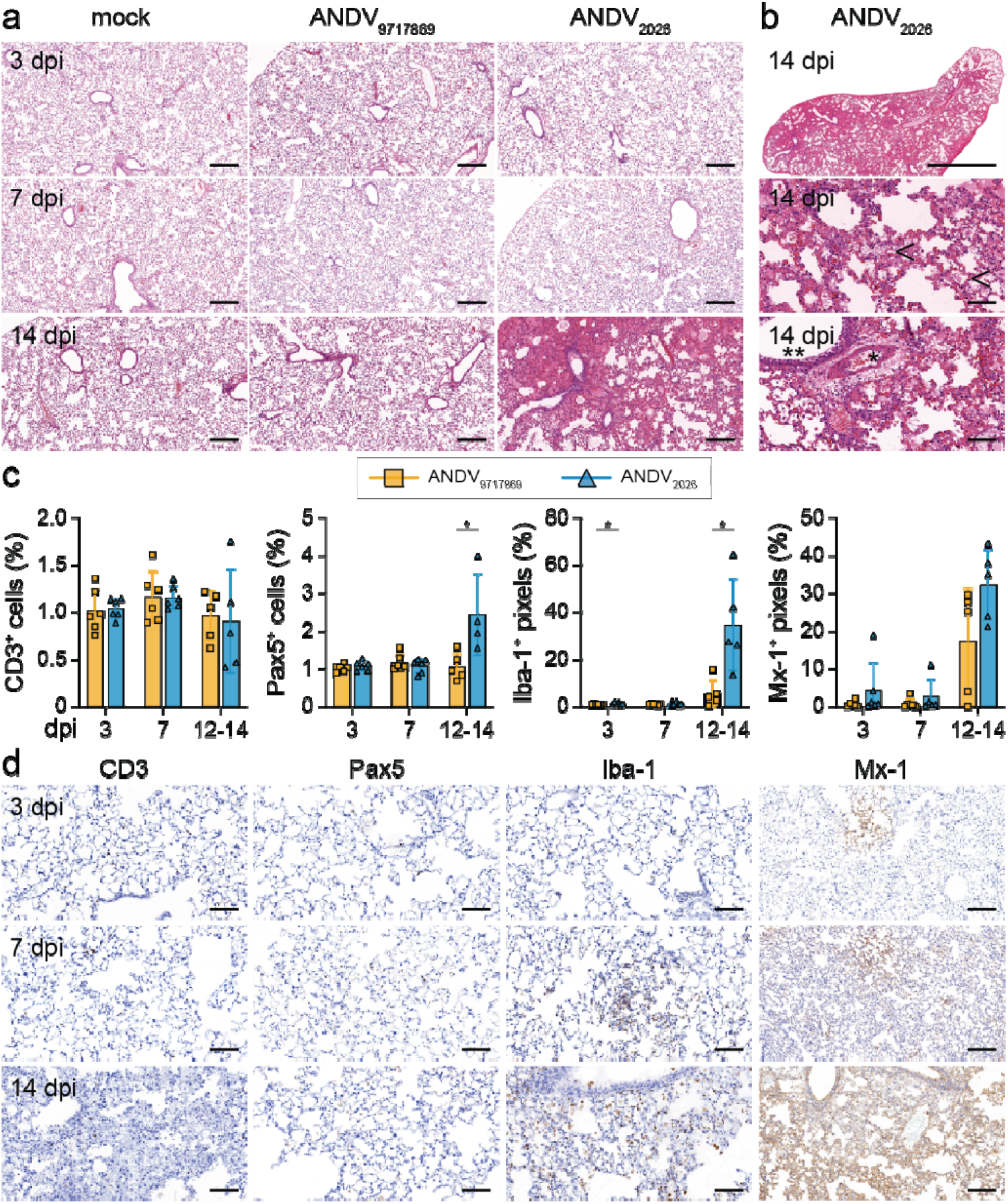
Severe pneumonia, macrophage infiltration and ISG upregulation in ANDV_2026_ inoculated hamsters. **(a)** Hematoxylin and eosin staining staining of FFPE lung sections of mock, ANDV_9717869_ and ANDV_2026_ inoculate hamsters at 3, 7 and 14 dpi. (**b**) Details of salient lung pathology at 14 dpi, top panel shows a lung lobe of a ANDV_2026_ inoculated hamster with lung oedema and a diffuse lesion distribution (bar, 3mm), accumulation of fibrin i the alveoli (arrow heads, middle panel) and mononuclear cell infiltrations in the widened perivascular space (*) as well as in the lung parenchyma; bronchus epithelium shows degenerated epithelial cells (**). Bar, 100µm. (**c**) Bar graphs representing software-assisted quantification of T cells (CD3), B cells (Pax5), macrophages (Iba-1), and Mx-1 positive signal at 3, 7 and 12-14 dpi in lungs of ANDV-infected animals as indicated. Data normalized on mock-infected animals (n=3). Data shown as mean ± SD; symbols represent individual hamsters. ANDV_9717869_ and ANDV_2026_ were compared by t-test or Mann-Whitney test for normal or non-normal distributed data, respectively; * p<0.05. (**d**) Representative images of immunohistochemistry of FFPE lung sections of ANDV_2026_ infected animals of T cells (CD3), B cells (Pax5), macrophages (Iba-1) and Mx-1 in mock, ANDV_9717869_ and ANDV_2026_ inoculated hamsters at 3, 7 and 14 dpi. Bar, 100 µm. Abbreviations: days post inoculation (dpi).

In addition, activation of the type I IFN response (henceforth IFN), assessed by the upregulation of the IFN-stimulated gene Mx-1, was evident in the lungs of both ANDV_2026_- and ANDV_9217869_-infected animals (**Fig. 3d**, last column). We detected relatively little Mx-1 at 7 dpi despite the presence of abundant virus replication. This observation reinforces the concept that ANDV is able to evade recognition by the host IFN response for some time.

Because ANDV, both in humans and in hamsters, causes primarily a pulmonary disease, we further investigated whether the levels of the IFN response vary between lungs and other organs (**Fig. 4**). To this end, we quantified the expression of IFIT-1 RNA, another interferon stimulated gene (ISG), in FFPE sections of lungs, spleen, myocardium, kidney and liver of ANDV_2026_-infected animals by RNAscope, and normalised the data in relation to the presence of viral RNA in each of these organs. We found the highest relative expression of IFIT-1 RNA in the lungs, followed by the spleen (**Fig. 4**). These data are in line with the relatively upregulation of proinflammatory genes such as CXCL10 that we detected in these two organs (**Fig. S3**). Of note, we did find bronchial epithelial cells expressing ISGs in the absence of viral replication (**Fig. 4a**), most likely triggered by a IFN response in infected neighbouring cells. The IFN and inflammatory response in respiratory epithelia may explain the lesions (sloughed off cells) observed in some of these animals.

**Figure 4.**
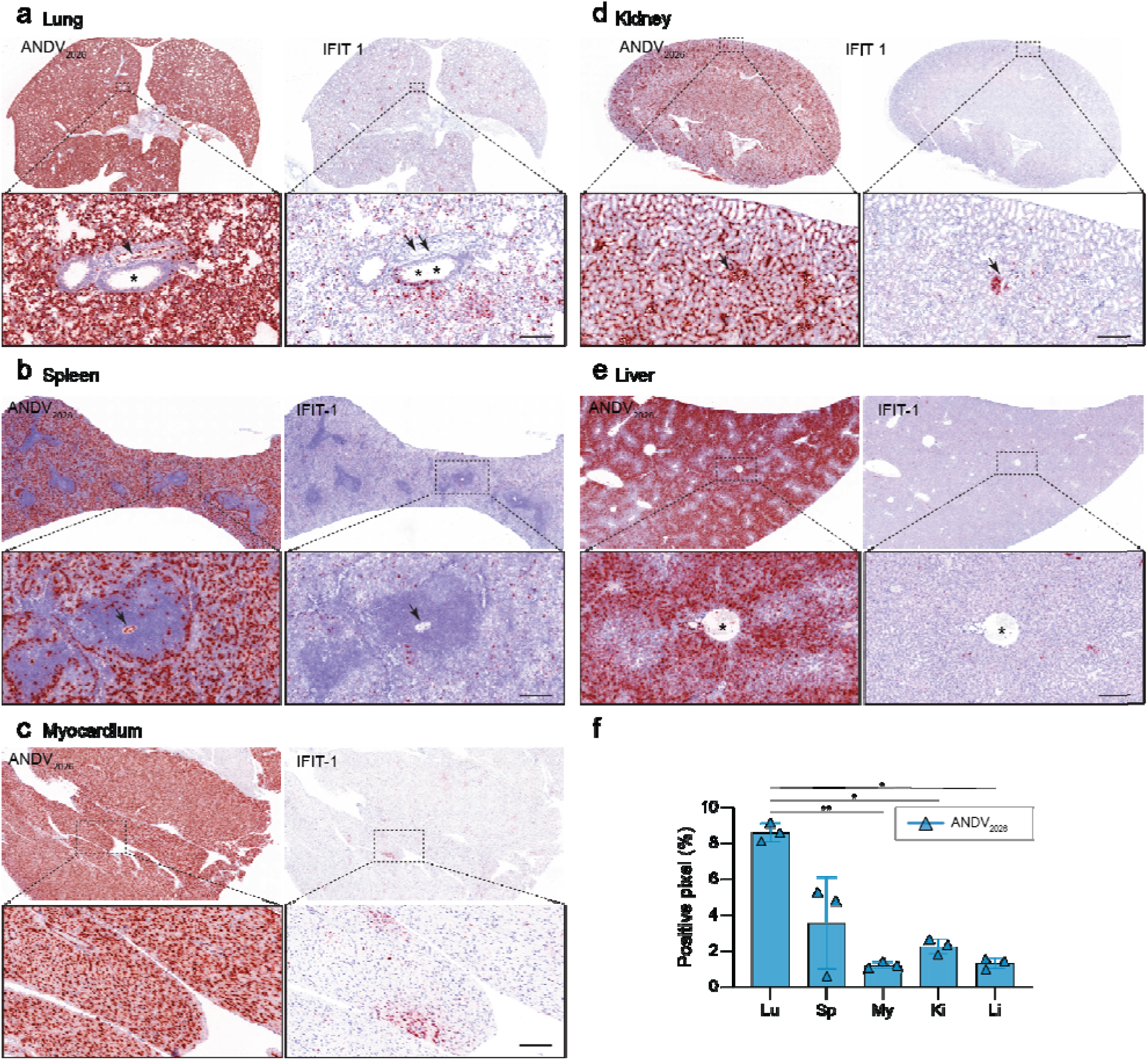
Detection of IFIT-1 in organs of ANDV_2026_ infected golden Syrian hamsters. Representative images of *in situ* hybridization detecting viral RNA of ANDV_2026_ and IFIT-1 at 12-14 dpi in lungs and peripheral organs. Consecutive serial sections are shown; the top row represents a low magnification whereas the bottom row represents a higher magnification of the indicated area of one hamster at 12 dpi. (**a**) Lung: * = virus-negative bronchus, arrow = virus-positive endothelial cells of a blood vessel; ** = IFIT-1 positive bronchial epithelial cells. Arrows = IFIT-1 negative blood vessel. (**b**) Spleen: arrow indicates a central artery surrounded by periarteriolar lymphatic sheaths (PALS). (**c**) Myocardium: staining pattern for the viral RNA is homogenous in the myocardium, while the staining for IFIT-1 shows patchy distribution. (**d**) Kidney: viral RNA is present in all glomeruli (arrow, left panel) and negative for IFIT-1 in the right panel (arrow), focal IFIT-1 signal in the tubuli below the arrow. (**e**) Liver: strong centro-lobular staining pattern of viral RNA and mild scattered signal for IFIT-1, central vein (*). All bars, 200µm. (**f**) Bar graphs of software-assisted quantification of IFIT-1 positive pixels signal, shown as percentage of ANDV_2026_ positive pixels. Data shown as mean ± SD; symbols represent individual hamsters. Data was compared b RM one-way ANOVA with multiple comparisons; * p<0.05, ** p<0.01. Abbreviations: lung (Lu); spleen (Sp); myocardium (My); kidney (Ki); liver (Li); days post inoculation (dpi).

Combined, ANDV_2026_ causes a disease comparable to HPS in hamsters, characterized predominantly by lung pathology, endothelial cell dysfunction, with a relatively long incubation period where abundant virus replication does not trigger a detectable inflammatory response.

### Adaptive immune response to ANDV in experimentally inoculated hamsters

We next assessed antibody and T-cell responses in ANDV-infected animals. We detected neutralizing antibodies only at 12-14 dpi in sera from animals infected with either ANDV_9717869_ or ANDV_2026_. The mean VNT_50_ did not differ significantly between groups, even though all ANDV_2026_-infected animals mounted a response compared to 40% of the ANDV_9717689_-infected animals (**Fig. 5a**). To further characterize the ANDV-specific adaptive immune response, we isolated lymphocytes from hamster lungs at both 7 and 14 dpi and stimulated them with overlapping peptide pools covering the N, Gn, Gc, and L proteins (the latter split in four distinct pools, L1-4) (**Fig. 5b**). At 7 dpi, antigen-specific T-cell responses were limited in all groups (**Fig. 5d**). However, strong antigen-specific T-cell responses were detected at 12-14 dpi regardless of group. The majority of virus-specific T-cells targeted the L-protein (**Fig. 5c, Fig. S4**); these were comparable between ANDV_9717869_- and ANDV_2026_-inoculated hamsters, and higher than those observed in mock-inoculated animals, although significance was reached only for ANDV_9717869_ (**Fig. 5e**). N-specific T-cell responses were higher in ANDV_9717869_-inoculated animals than in ANDV_2026_- and mock-inoculated animals at 12-14 dpi (**Fig. 5e**). Similar responses were observed with lymphocytes isolated from the spleen, although the background in mock-inoculated hamsters was higher, complicating data interpretation (**Fig. S4**).

**Figure 5.**
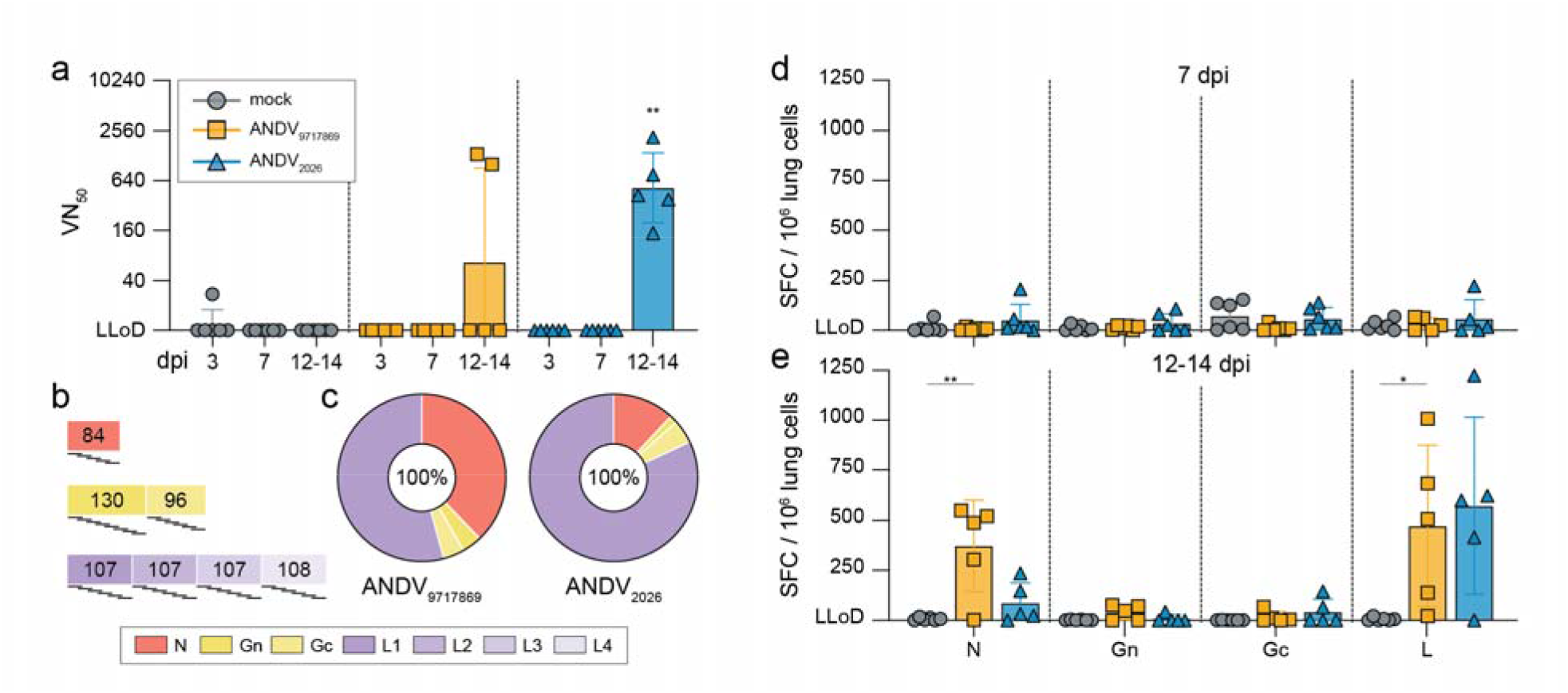
ANDV-specific antibody and T cell responses in experimentally inoculated hamsters. (**a**) ANDV_9717869_ neutralizing antibody levels measured at 3, 7 and 12-14 dpi. Neutralizing antibody levels are expressed as VN_50_, data shown as geometric mean ± geometric SD interval; symbols represent individual hamsters. Log-transformed ANDV_9717869_ and ANDV_2026_ were compared to mock by Kruskal-Wallis with multiple comparisons for non-normal distributed data; ** p<0.01. (**b**) Schematic representation of peptide pools used to stimulate lymphocytes for the detection of virus-specific T cell responses. L was covered by 4 peptide pools (L1-4). Numbers represent the number of peptides per pool. (**c**) T cell response specificity shown as fraction of the total T cell response, based on absolute data shown in panel e. (**d, e**) ANDV-specific T cell responses in the lungs measured by stimulation of lymphocytes with overlapping peptide pools and IFNg ELISPOT at 7 and 12-14 dpi. T cell responses are expressed as spot forming cells (SFC) per 10^6^ total cells. T cell responses targeting L are shown as the sum of L1-, L2-, L3-, and L4-specific responses. Data shown as mean ± SD; symbols represent individual hamsters. Log-transformed ANDV9717869 and ANDV2026 data were compared to mock by Kruskal-Wallis or one-way ANOVA with multiple comparisons for non-normal or normal distributed data, respectively; * p<0.05, ** p<0.01. Abbreviations: lower limit of detection (LLoD); virus neutralization (VN); nucleoprotein (N); glycoprotein N (Gn); glycoprotein C (Gc); polymerase (L); days post inoculation (dpi).

### Infectious virus is shed in the air by ANDV_2026_-infected animals

The ANDV outbreak aboard the *MV Hondius* raised questions about the potential of increased shedding, and hence increased transmissibility, of ANDV. In hamsters, we detected viral genomes (**Fig. 6a**) and infectious virus (**Fig. S5**) in throat swabs of all ANDV_2026_-inoculated animals from 7 dpi onwards, with progressively increasing titers until the end of the experiment (**Fig. 6a**). We detected infectious virus in only one ANDV_9717869_-infected animal. By immunohistochemistry, we detected numerous clusters of ANDV_2026_ infected endothelial cells, olfactory epithelium, degenerated and sloughing olfactory epithelium, as well as the ducts of the nasal submucosal glands of the olfactory portion of the turbinates, especially at 14 dpi. Only very few cells of the respiratory epithelium showed occasionally virus positive signal (**Fig. 6d-f**). ANDV_9717869_-inoculated animals displayed only occasional virus-positive cells in the nasal turbinates (either endothelial cells, cells within the olfactory epithelium, or the submucosal glands).

**Figure 6.**
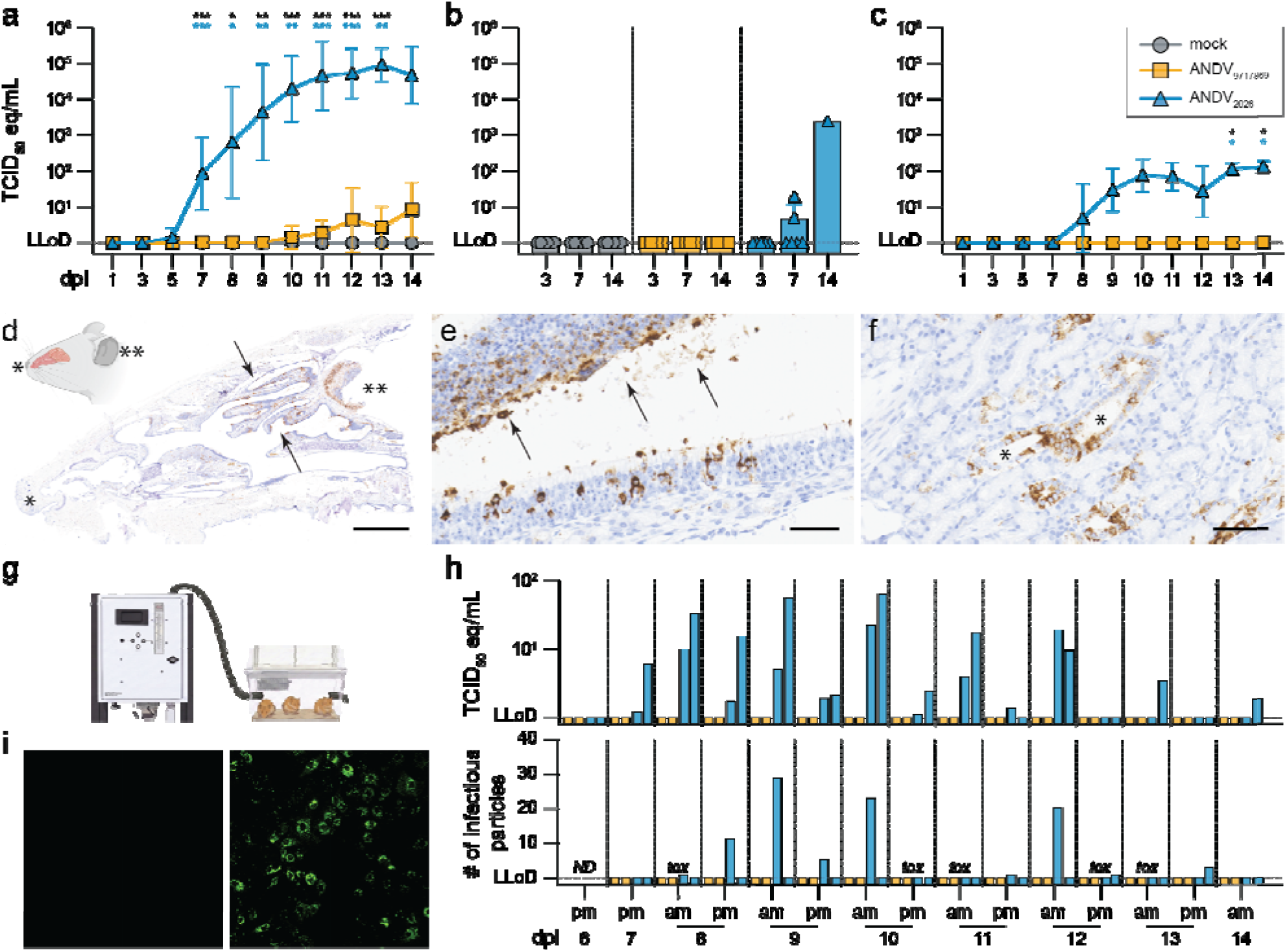
ANDV shedding in experimentally inoculated hamsters. (**a**) Quantification of viral genome in throat swabs of mock, ANDV_9717869_ and ANDV_2026_ inoculated hamsters measured by qRT-PCR. Data shown as geometric mean ± geometric SD. Lines were compared by a two-way ANOVA mixed effects analysis with multiple comparisons on log-transformed data. Colour-coded asterisks denote the difference between respective group and mock, black asterisks between two ANDV groups; * p<0.05, ** p<0.01, *** p<0.001. (**b**) Quantification of viral genome in urine, conditionally obtained during necropsies of mock, ANDV_9717869_ and ANDV_2026_ inoculated hamsters measured by qRT-PCR. Data shown as geometric mean ± geometric SD; symbols represent individual hamsters. (**c**) Quantification of viral genome collective fecal homogenates of mock, ANDV_9717869_ and ANDV_2026_ inoculated hamsters measured by qRT-PCR. Data shown as geometric mean ± geometric SD. Lines were compared by a two-way ANOVA mixed effects analysis with multiple comparisons on log-transformed data. Colour-coded asterisks denote the differenc between respective group and mock, black asterisks between two ANDV groups; * p<0.05. (**d**) Overview of nasal turbinates showing abundant viral protein between the arrows (IHC); tip of the nose (*), and brain cavity for orientation (**). Bar, 3 mm, 14dpi. (**e**) High magnification of olfactory epithelium with virus-positive olfactory epithelium with maintained morphology (bottom) and degenerated cells with epithelial sloughing (top, arrows). Bar, 50 µm, 14 dpi. (f) Virus-positive duct epithelium (*) of the nasal submucosal glands. bar, 70 µm, 14 dpi. (**g**) Schematic representation of airsampling setup including hamster cage and BioSpot-VIVAS. (**h**) Quantification of ANDV genome equivalents (top panel), expressed as TCID₅₀ equivalents/mL, and infectious virus (bottom panel), quantified by virus isolation in Vero E6 cells, in air samples collected from ANDV9_717869_- and ANDV_2026_-inoculated hamsters. Bars show levels in individual airsamples collected either in the morning (am) or evening (pm). ND indicates timepoint on which isolation was not performed, tox indicates timepoints on which airsample toxicity interfered with detection of ANDV infected cells. (**i**) Representative staining of ANDV_2026_ isolated from airsamples on VeroE6 cells (right panel). Isolation from a sample without detectable ANDV_2026_ is shown as a negative control (left panel). Images were taken at a 200x magnification. Abbreviations: days post inoculation (dpi).

We also detected ANDV_2026_ RNA in urine collected at study endpoints, but volumes were too small to test for presence of infectious virus. Furthermore, we detected both viral RNA and infectious virus in feces homogenates collected from cages housing ANDV_2026_ infected animals (**Fig. 6c and S5**).

Hence, ANDV_2026_-infected animals shed infectious virus from the respiratory tract, urine and feces. To further investigate whether ANDV_2026_ was aerosolized, we connected the cages housing infected hamsters to a BioSpot-VIVAS series 315 bioaerosol sampler. Air was sampled continuously in 12-hour increments (**Fig. 6g**). Beginning at 7 dpi, ANDV RNA was detected in air samples collected from cages housing hamsters that were inoculated with ANDV_2026_ but not from cages housing animals inoculated with ANDV_9717869_ **(Fig. 6i**). Shed viral RNA peaked around 9 and 10 dpi (**Fig. 6h**). Infectious virus was detected in most air samples containing viral RNA, though only from cages of hamsters infected with ANDV_2026_ (**Fig. 6h**). Due to the cytotoxic effects of the collected air samples on the cells used for titration, infectious viral particle counts are likely underestimated (**Fig. 6h**). Overall, ANDV_2026_ is shed via multiple routes by infected hamsters and could be detected in aerosol samples as both viral RNA and infectious virus.

## DISCUSSION

Here, we performed a comprehensive *in vivo* characterisation of the recently emerged ANDV isolate from the *MV Hondius* cruise ship outbreak. To assess how this isolate differed from previously described isolates, and to provide a comprehensive risk assessment, we evaluated its pathogenicity, the virus-induced innate and adaptive host responses, as well as its shedding potential in golden Syrian hamsters, the animal model for HPS. ANDV_2026_ proved to be highly virulent, causing in hamsters a disease similar to HPS in humans, characterized primarily by pulmonary pathology. This was in contrast to previous studies that used human-derived ANDV isolates, usually displaying an attenuated phenotype^22,24^.

ANDV_2026_ spread systemically to multiple organs by targeting endothelial cells. Indeed, we detected ANDV_2026_ in every organ that we tested, including among others spleen, myocardium, kidneys, urinary bladder, thymus, adrenal gland and intestine as also shown in previous studies^22,24,29^. We also found extensive ANDV_2026_ infection in the brain, including in blood vessels, meninges, and neurons, previously unrecognized targets of ANDV replication. Together with reports of ANDV-associated neurological manifestations, our data indicate that this virus is potentially neuroinvasive^30^.

Despite systemic spread of ANDV, macroscopic and histopathological lesions were largely confined to the lungs. Disease severity was driven mainly by alveolar damage, accompanied by alveolar and perivascular oedema and in some cases sloughing of the bronchial epithelium. Using software-assisted image analysis, we localised viral antigen and RNA to the lung parenchyma, with no detectable signal in the bronchial epithelium. The IFN response, as visualised by expression of ISGs such as Mx-1 and IFIT1, also extended to bronchial epithelial cells. Therefore, pathology correlated with preferential IFN activation in the lungs relative to other organs, including myocardium, kidney and liver, with macrophages infiltrate, in addition to increases in pulmonary inflammatory markers. The latter have been reported also in previous studies with ANDV in hamsters^26,29,31^ and other hantaviruses in humans^32,33^. These data are in accord with observations *in vitro* in human cells suggesting that infected endothelial cells may coordinate leukocyte recruitment rather than be cytokine-producing effector cells^34^. More studies are needed to determine mechanistically why the extent of the IFN response in ANDV-infected endothelial cells is lower than in the lung parenchyma. It is also important to note that even in the lungs, where ANDV replicates abundantly at 7 dpi, the IFN response is relatively modest compared with, for example, what we previously detected in hamsters infected with acute respiratory viruses such as SARS-CoV-2^28^. These data are consistent with the many immunoevasive properties of ANDV described in *in vitro* systems^35,36^. We also argue that IFN induction via damage associated molecular patterns (DAMPs) will be reduced compared to other acute respiratory infections, considering the ANDV is not cytopathogenic. Self DNA and RNA is increasingly being recognised as key triggers of the IFN response^37–41^.

In addition to the innate immune responses, we characterized adaptive immune responses in infected hamsters. As in previous studies^25^, we detected neutralizing in the late stages of the disease, but they did not facilitate viral clearance. Although it has been hypothesized that T-cells could play a role in the immunopathogenesis HPS^42^, a previous study showed that disease development in hamsters is not influenced by either CD4+ or CD8+ T cells^35,36^. We did not detect an increase in T-cells in the lungs of ANDV-infected animals. We detected however antigen-specific T-cell responses in lungs and spleens at 7 and 12-14 dpi in both ANDV_2026_- and ANDV_9717869_-inoculated animals, targeting predominantly epitopes in the L protein. Additionally, T-cell reactivity towards the N protein was mainly observed between in ANDV_9717869_-inoculated animals, potentially due to a preferential affinity of epitopes to specific MHC molecules. In any case, we found no correlation between T-cell responses and protection from clinical signs of ANDV infection, similarly also to what observed in infected patients^43^.

Overall, our data on the adapted host responses to ANDV leads to an interesting discordance: neutralizing antibodies target the viral Gn/Gc, whereas the N and L protein seem to be the immunodominant target of the T-cell response. This is, to our knowledge, the first report of differential T-cell reactivity towards the entire viral proteome of ANDV and it may provide important perspectives for future vaccine strategies.

Lastly, we found ANDV_2026_ to be shed by infected animals through various routes and, importantly, to be aerosolized. Data related to ANDV shedding is limited in humans^16^. Infectious virus has been detected in urine^44^ and viral RNA in saliva, gingival crevicular fluid, and nasopharyngeal swabs^45^, with isolated reports in semen and breast milk^46,47^.

We detected infectious virus in throat swabs from 7 dpi onwards, in addition to feces and urine in the ANDV_2026_-infected group. Virus detection remained limited in throat swabs and absent in the feces and urine of ANDV_9717869_-inoculated animals. Also in previous reports, the shedding potential of ANDV_9717869_ was shown to be limited^48^ or absent^27^, highlighting a defining phenotype for ANDV_2026_. We found abundant virus antigen in the olfactory epithelium and endothelial cells in the nose of ANDV_2026_-infected animals. Virus positive cells in the nasal turbinates of ANDV_9717869_ were sporadically found, in line with previous studies in hamsters^26^. Indeed, we showed that titers of infectious virus that we recovered from turbinates of ANDV_2026_-infected animals were about 100-fold higher at 12-14 dpi than those recovered from the nasal turbinates of ANDV_9717869_-infected animals. This may provide a route of entry into the bloodstream, considering that the bronchial epithelium was virus negative throughout the duration of the experiment.

Beyond shedding via multiple routes, we also recovered infectious virus from the air of cages housing ANDV_2026_-inoculated hamsters. To our knowledge, this is the first isolation of an orthohantavirus from air, representing an important addition to the evaluation of its transmission potential. Whether these aerosolized particles are sufficient to support transmission remains to be determined. By contrast, air sampling in a hospital during the current outbreak did not detect virus^49^. Although direct comparisons between the two settings should be made with caution, the negative hospital findings may reflect differences in ventilation, air exchange rates, sampling conditions, or levels of virus shedding between infected patients and experimentally infected hamsters. The actual transmission potential of ANDV will require dedicated follow-up studies assessing transmission to immunologically naïve contact animals. Nonetheless, our data strongly indicate efficient shedding of ANDV_2026_ via multiple routes.

A limitation of our study is that the phenotype of ANDV_9717869_ in hamsters differs from that reported elsewhere, where it usually caused lethal infection^25–27^. This discrepancy may reflect the low challenge dose used here: 89 TCID_50_ for ANDV_9717869_ (and 28 TCID_50_ for ANDV_2026_), based on back titration of the inocula. The LD_50_ for intranasal ANDV inoculation has been estimated at approximately 95 PFU^50^. It is therefore even more striking that ANDV_2026_ caused such severe disease, underscoring the substantial pathogenic potential of this isolate.

Overall, the *MV Hondius* hantavirus outbreak provides a stark reminder of the impact that emerging infectious diseases can have globally, even when affecting a relatively low number of patients from a public health perspective. Early phenotypic characterization of emerging viruses is a critical step of any outbreak response as it facilitates a comprehensive risk assessment and provides the foundation for development and assessment of tailored countermeasures.

## Methods

### Ethics statement

Research involving animals was conducted in compliance with the Dutch legislation for the protection of animals used for scientific purposes (2014, implementing EU Directive 2010/63) and other relevant regulations. The licensed establishment where this research was conducted (Erasmus MC) has an approved Office of Laboratory Animal Welfare (OLAW) Assurance # F16-00046 (A5051-01). Research was conducted under a project license (#2619213) from the Dutch competent authority, and the study protocol was approved by the institutional Animal Welfare Body (Erasmus MC). Animals were housed in groups of 3 animals in filter top cages (Type 3, Techniplast), in Class III isolators, allowing social interactions, under controlled conditions of humidity, temperature and light (12-h light/12-h dark cycles). Food and water were available ad libitum. Animals were cared for and monitored daily by qualified personnel. All animals were allowed to acclimatize to husbandry for at least 7 days. For unbiased experiments, all animals were randomly assigned to experimental groups. The animals were anesthetized (3% isoflurane) for all invasive procedures. Hamsters were euthanized by cardiac puncture under isoflurane anesthesia.

### Cell culture

Vero E6 cells (#CRL-1586, ATCC) were cultured in Dulbecco’s Modified Eagle Medium (DMEM; #DMEM-HPXA, Capricorn Scientific) supplemented with 10% fetal bovine serum (FBS; #F7524, Sigma), 2 mM L-glutamine (#GLN-B, Capricorn Scientific), 0.5 µg/mL amphotericin B (Bristol Myers Squibb), and 100 units/mL penicillin and 0.1 mg/mL streptomycin (#PS-B, Capricorn Scientific) at 37 °C and 5% CO_2_ in a humidified incubator.

### Viruses

ANDV/2026/hondius-NL-12 (ANDV_2026_) was isolated in VeroE6 cells from a whole-blood sample (PCR-positive for ANDV, pre-seroconversion), which was collected from a pre-symptomatic ANDV patient during the 2026 ANDV outbreak linked to the cruise ship *MV Hondius*^3^. ANDV 9717869/AF291704 (ANDV_9717869_) was acquired from the European Virus Archive Global (#002v-EVA400). Virus stocks were propagated on VeroE6 cells. Additionally, ANDV_9717869_ stocks were concentrated using ultrafiltration. Briefly, supernatant from ANDV_9717869_-infected VeroE6 cells was collected, clarified (5,000*g*, 10 min, 4 °C), and ultrafiltrated (4,000*g*, 40 min, 4°C using Amicon® Ultra Centrifugal Filter, 100 kDa MWCO [#UFC910024; Merck Millipore]), before eluting concentrated ANDV_9717869_ from the filter membranes.

### Animals and experimental setup

Female and male golden Syrian hamsters (n=54; *Mesocricetus auratus*; 6 weeks old; Janvier, France) were handled in an ABSL-3 biocontainment laboratory. Animals were intranasally inoculated with either 200 TCID_50_ of ANDV_2026_ (n=18) or ANDV_9717869_ (n=18) or sterile PBS (n=18) in a total volume of 50 µl per animal. The diluted inocula were backtitrated to further control for the infectious titer; resulting titers were 28 TCID_50_/50 µl for ANDV_2026_ and 89 TCID_50_/50 µl for ANDV_9717869_. Two animals had to be excluded from the experiment pre-terminally due reasons irrelevant to the experiment. Consequently, the group size at 12-14 dpi is decreased (n=5).

In addition, to confirm that animals were inoculated with the intended isolates, viral RNA collected from nasal turbinates of animals from groups killed at 3 dpi were also sequenced (**FigS6, TabS1**). As expected, all samples confirmed the intended experimental setup (see material and methods below). Throat swabs and collective feces samples per cage were taken at 1, 3, 5, 7, 8, 9, 10, 11, 12, 13 and 14 days post inoculation (dpi). The body weight of animals was measured daily. From 7 dpi onwards, two cages of animals inoculated with either ANDV_2026_ or ANDV_9717869_ were connected to air sampling devices. On 3 dpi, 7 dpi, and 14 dpi, animals (nL=L6) from each experimental group were euthanized and tissues (nasal turbinates, brain, submandibular glands, cervical lymph node, trachea, lung, heart, thymus, liver, pancreas, spleen, kidney, adrenal gland, small intestine and urinary bladder) as well as urine, if found in the urinary bladder, were collected for quantification of viral and genomic load, as well as for histopathology and virus antigen and RNA expression. Lungs and spleens were furthermore subjected to lymphocyte isolation, and the weight of the lungs was recorded.

### Titration of virus stocks and samples from ANDV-infected hamsters

The titers of virus stocks and samples (throat swabs, fecal homogenates, tissues) from ANDV-infected hamsters were quantified by endpoint dilution (TCID_50_ assay, 50% tissue culture infectious dose). To this end, tenfold serial dilutions in infection medium (Minimum Essential Medium [#SP-2202-500ML, Capricorn Scientific] supplemented with 10% FBS, 2.2 g/L sodium bicarbonate [#25080060, Gibco], 10 mM HEPES [#15630056, Gibco], 2 mM L-glutamine, 0.5 µg/mL amphotericin B, and 100 units/mL penicillin and 0.1 mg/mL streptomycin) were prepared and 100 µL of each dilution to were transferred to 70%-90% confluent VeroE6 cells in 96-well cell culture plates. The cells were incubated for 5 days at 37 °C and 5% CO_2_ before fixation for 30 min using 10% neutral buffered formalin and permeabilization in 70% ethanol by submerging the entire plate. To visualize ANDV-infected cells, they were washed three times with PBS, blocked in blocking buffer (0.6% BSA [#A8327, Sigma-Aldrich] and 0.1% Triton X-100 [#T8787, Sigma-Aldrich] in PBS), and stained for 2 h at room temperature with the monoclonal TULV 1 antibody (1:50-1:100; kindly provided by Sven Reiche, Friedrich-Loeffler-Institut, Germany)^51^. To determine the titer of the two virus stocks as well as to quantify the back titration of the input of the hamster inoculation, the monoclonal anti-hantavirus nucleocapsid protein antibody 5E11 (1:100; #ab34757, abcam) was used. After washing with PBS, cells were incubated with an Alexa Fluor 488-conjugated goat anti-mouse antibody (1:500; #A-11001, Invitrogen) or donkey anti-mouse antibody (1:500; #A-21202, Invitrogen). Following another wash step with PBS, infection was assessed using a fluorescence microscope (Zeiss Axio Vert.A1). Viral titers were calculated using the Spearman-Kärber method and expressed as TCID_50_/mL or TCID_50_/g.

### RNA isolation and ANDV PCR

All throat swabs, feces homogenates, air samples and tissue samples were used for RNA extraction and following quantification of ANDV genome by RT-qPCR. RNA extraction was performed as described previously^52^. A primer-probe set (HCPS fwd 1: ATGACGTTAACAARAGCACATTACA [9.6 pmol/reaction]; HCPS-fwd_2: ATGACGTTAACAARAGCACACTACA [9.6 pmol/ul]; HCPS_rev: TCAAKCCCTGTTGGATCAA [14.4 pmol/ul]; HCPS_probe: FAM-TCCAATGYAGACACAGCTGCCCGT-BHQ1 [0.8 pmol/ul]; HCPS_probe_2: FAM-TCCAGTGTAGACACAGCTGCCCTT-BHQ1 [0.8 pmol/ul]) targeting the S-segment was utilized (van Beek et al., 2026, submitted). Genome was subsequently amplified (50°C, 5min; 95°C, 20sec; 45 cycles of 95°C, 3 sec and 60°C, 31 sec). Ct values were compared to a standard curve derived from a titrated ANDV stock. Host genes (IL6, IL1ß, TNFα, IFNγ and CXCL10) were measured utilizing hamster specific primers and probes^53^ and were normalized to the housekeeping gene ß-actin in a duplex RT-qPCR for all lung and spleen samples.

### Sequencing of ANDV

ANDV whole-genome nanopore sequencing was performed on virus inoculum and nasal turbinate tissue collected at 3 days post inoculation. Turbinate tissues were initially homogenized in virus transport medium and centrifuged for 5 minutes at 10,000 × g. A total of 200 μL of supernatant was added to a mixture of 200 U OmniCleave Endonuclease (Biosearch Technologies), 25 mM Tris-HCl pH 8.0 and 4 mM MgCl2, followed by incubation for 30 minutes at 37°C to digest free nucleic acids. Total nucleic acids were then isolated using the High Pure Viral RNA isolation kit (Qiagen) according to manufacturer’s instructions. Following cDNA synthesis an amplicon-based sequencing assay, developed specific for the 2026 ANDV which caused the outbreak on a cruise ship, was applied for whole-genome sequencing on a GridION machine (Oxford Nanopore Technologies) as described in van Beek et al., 2026 (manuscript in revision). Data analysis was performed as previously described in (van Beek et al., 2026, manuscript in revision), using a minimum read depth of 20× for consensus calling.

### Airsampling

Cages housing inoculated hamsters were connected to BioSpot-VIVAS series 315 bioaerosol samplers (Aerosol Devices), using a 1.4-meter long tubing. BioSpots were operated at a flow rate of 15L/min, under the following conditions: conditioner: 5 °C; initiator: 45L°C; moderator: 18 °C; nozzle: 27 °C; sample holder: between 10 °C and 17 °C, as described previously^54^. With a cage volume of 24 L and a flow rate of 900LL/hour (60L×L15LL/min), this results in 37,5 air changes per hour. The hamster cages and BioSpot were housed in separate, interconnected, negatively pressurized class 3 isolators. This separation was necessary because the BioSpot generated excessive heat, which led to an increased ambient temperature within the isolator. Sampled aerosols were collected in 2 mL of MEM with Hanks salt solution (Gibco), supplemented with 5 mg/mL lactalbumine enzymatic hydrolysate (Sigma-Aldrich), 200 IU/mL penicillin and 200 ug/mL streptomycin, 100 U/mL Polymyxine B (Sigma-Aldrich), 5 MU Nystatin (Gibco), 250 µg/mL gentamycin (Invitrogen), and 10% Glycerol (Sigma-Aldrich) (VTM). Air sampling was performed continuously from 7 to 14 dpi, with sample collection every 12 hours. Bedding material in the cages was changed every morning to prevent the accumulation of virus in the bedding, which could influence air samples. Collected air samples were immediately adjusted to a final volume of 5 ml with infection medium, and used to inoculate half of a 96-well plate containing 70%-90% confluent VeroE6 (100 µL/well). Inoculated cells were incubated at 37 °C, 5% CO_2_, for 5 days, after which plates were fixed with 10% neutral-buffered formalin for 30 minutes, followed by washing with 70% ethanol. Immunostaining of ANDV-infected cells was done as described earlier.

### Lymphocyte isolation from spleen and lung

Spleens were collected during necropsy in Iscove’s Modified Dulbecco’s Medium (IMDM, Capricorn Scientific) supplemented with 10% FBS (I10F) and primocin (100 µg/ml, Invivogen) pending processing. Single cell suspensions were manually generated by straining through a 100 µm cell strainer (Falcon). Cells were subsequently washed with Roswell Park Memorial Institute 1640 (RPMI, Capricorn Scientific) supplemented with 10% FBS (R10F) and primocin (100 µg/ml) and incubated in red blood cell lysis buffer (Roche) for 10 minutes. After lysis, single cell suspensions were washed twice with R10F supplemented with primocin (100 µg/ml) and kept on ice pending analysis. Lungs were collected during necropsy in I10F supplemented with primocin (100 µg/ml) pending processing. Lung tissues were manually processed into 0.5-1 mm^3^ pieces and subsequently homogenized in I10F supplemented with collagenase D/IV (1 mg/ml, Worthington), DNase I (100 ug/ml, Worthington), soybean trypsin inhibitor (1 mg/ml, Thermo Fisher), sodium pyruvate (1 mM, Gibco), non-essential amino acids (1x, Capricorn Scientific), penicillin/streptomycin (100 IU/100 µg/ml,), L-glutamine (2LmM,), and primocin (100 µg/ml) using the GentleMACS system (Miltenyi Biotec). Cell homogenates were subsequently incubated at 37 °C for one hour and further homogenized. Single cell suspensions were generated by straining through a 100 µm cell strainer (Falcon), followed by washing with I10F supplemented with primocin (100 µg/ml). Subsequently, cells were incubated with DNase I (50 µg/ml) in I10F supplemented with primocin (100 µg/ml) at 37LC for 15 minutes, treated cells were washed and finally loaded onto a density gradient for lymphocyte isolation (Percoll®, Cytiva). Lymphocytes collected from the interphase were washed twice and kept on ice pending analysis.

### Peptide synthesis and megapool preparation

Overlapping peptide pools (15-mers with 10 amino acid overlap) that span the nucleoprotein (N), glycoprotein (Gn and Gc), and polymerase (L) open reading frames were designed and produced in-house. Gc and Gn pools were based on ANDV/2026/hondius-NL-12 (Pathoplexus accession # PP_006W3U9.2), N and L pools were based on ANDV_9717869_ (NCBI accession # NP_604471.1 and NP_604473.1, respectively). Peptides were synthesized as crude material (TC peptide lab) and individually resuspended in dimethyl sulfoxide (DMSO). Aliquots of all peptides were pooled and sequentially lyophilized into megapools as previously described^55^, designated as ANDV-N, ADNV-Gn, ADNV-Gc, ANDV-L-P1, ANDV-L-P2, ANDV-L-P3, and ANDV-L-P4. Each individual peptide was present at a concentration of 1 mg/mL in the megapool stock solution, used at a final test concentration of 1 µg/mL per peptide. All peptides and megapools are listed in **Table S2**.

### IFNγ ELISpot

Hamster IFNγ enzyme-linked immunospot (ELISpot, Mabtech) was performed according to the manufacturer’s instructions. Briefly, plates were conditioned with R10F or I10F in preparation of stimulation of splenocytes or lymphocytes isolated from the lungs, respectively. Subsequently, plates were loaded with peptide pools (ANDV-N, ADNV-Gn, ADNV-Gc, ANDV-L-P1, ANDV-L-P2, ANDV-L-P3, and ANDV-L-P4, 1 µg/ml) and cells were added (5x10^5^ cells/well). An equimolar concentration of DMSO was included as negative control; phorbol 12-myristate 13-acetate (PMA) and ionomycin was included as positive control. ELISpot plates were incubated at 37LC and 5% CO2 for 36 h. Plates were washed five times using PBS and stained with a biotinylated anti-hamster IFNγ monoclonal antibody (Mabtech) for two hours at room temperature, washed five times with PBS, and incubated with streptavidin-HRP (Mabtech) for 1 h at room temperature. Plates were washed and subsequently developed using TMB substrate (Mabtech) for 10 minutes. Spot forming cells (SFC) were counted using a CTL Immunospot with pre-defined settings for either splenocytes or lymphocytes isolated from the lungs. Total SFC were corrected for background by subtracting DMSO-stimulated wells and visualized as SFC per 1 million cells.

### Histology, immunohistochemistry and *in situ*-hybridisation

2-3 μm thick sections of formalin-fixed and paraffin-wax embedded (FFPE) tissues (lung, spleen, liver, myocardium, kidney, pancreas, thymus, small intestine, brain, submandibular glands and lymph nodes, adrenal gland, urinary bladder, skeletal muscles, and EDTA-decalcified turbinates) were cut and mounted on charged glass slides using standard histological procedures. Slides were stained with haematoxylin and eosin (HE). For immunohistochemistry, the following antibodies were used: anti-TULV-1 (provided by Sven Reiche, Friedrich-Loeffler-Institut, Insel Riems, Germany), anti-N Andes 1A8-F6 (Amsbio), anti-CD3 (Agilent DAKO) anti-Pax5 (Agilent DAKO), anti-Iba-1 (Fujifilm), and anti-Mx-1 (Cell Signaling). As negative controls, we used mouse monocolonal isotype control antibodies or normal rabbit serum. For visualization, the EnVision+/HRP, Mouse, HRP kit (Agilent DAKO) or EnVision+/HRP, Rabbit, HRP kit (Agilent DAKO), respectively, was used in an automated stainer (Leica Bond RX, Autostainer Link 48, Agilent Technologies). For all immunohistochemistry experiments, 3,3’-Diaminobenzidine (DAB) was used as a chromogen. RNA was detected using RNAScope according to manufacturer’s instructions (Advanced Cell Diagnostics, Bio-Techne Spatial) with simmering in target-retrieval solution and proteinase K treatment. The following probes were used: ANDV_9717869_ (# 900241), ANDV_2026_ (#1908411), DapB (#310043), Ubiquitin (#310041), as well as IFIT-1 (#1153111). Counterstaining was performed with haematoxylin.

### Signal quantification by digital pathology

All slides were digitized with an Aperio VERSA 8 Brightfield, Fluorescence & FISH Digital Pathology Scanner (Leica Biosystems) at 200 x brightfield magnification or with the Zeiss Axioscan 7 using the Plan-Apochromat 20x/0.8 objective.

To quantify the positive signal of Fast-red (*in situ*-hybridisation, ISH) and diamino-benzidine (DAB)-brown (immunohistochemistry, IHC) in the tissues (nasal turbinates, lung, kidney, liver, spleen and myocardium), we optimised image analysis pipelines based on the signal detected using the QuPath software (version 0.7.0; Bankhead et al.)^28,56^. The pixel classifier (Iba-1, Mx-1, viral proteins, viral RNA) or cell detection (CD3, Pax-5) tools have been optimized for each staining separately. All slides with a specific antibody or probe have been analysed with the same settings after annotating the region of interest. Halo version 4.1.5944 (Indica labs, Inc.) was used to determine the virus-positive signal in the bronchus and parenchyma, respectively. After training HALO and HALO AI with the DenseNet V2 classifier, the software was able to distinguish between a) virus-positive bronchus, b) virus-negative bronchus, c) virus-positive parenchyma and d) virus-negative parenchyma and calculated the respective areas.

### Detection of ANDV-neutralizing antibodies

To determine the level of ANDV-neutralizing antibodies, we adapted an established microneutralization protocol^57^. To this end, VeroE6 cells were seeded one day prior in 96-well cell culture plates to yield 70%-90% confluent monolayers the following day. Twofold serial dilutions in infection medium of heat-inactivated sera (30 min, 56 °C; 1:10 starting dilution) were added to an equal volume of infection medium containing 200 TCID_50_ ANDV_9717869_ or ANDV_2026_. All measurements were performed in duplicate. Following a 2 h incubation of the virus-serum mix at 37 °C, 100 µL were transferred to 96-well cell culture plates, which were incubated for 5 days at 37 °C and 5% CO_2_. Two h post-inoculation, all wells were topped up with an additional 100 µL infection medium. Following incubation, cells were fixed using 10% neutral-buffered formalin for 30 min and permeabilized in 70% ethanol by submerging the entire plate. Immunostaining of ANDV-infected cells was done as described earlier. Neutralizing titers were determined by assessing total infected area per well using virus-specific fluorescence using a CTL Immunospot. VNT_50_ was defined as the highest dilution of serum at which at least 50% of the infected area was reduced compared to the virus control. When no neutralization was observed, the VNT_50_ was assigned a value of 10.

### Clinical chemistry

Sera of hamsters sacrificed on 12-14 dpi were subjected to an evaluation of clinical chemistry values. Samples were tested on the VetScan VS.2 clinical chemistry analyzer (Zoetis diagnostics) with the comprehensive diagnostic profile, testing albumin, alkaline phosphatase, alanine aminotransferase, amylase, blood urea nitrogen, calcium, creatinine, globulin, glucose, potassium, sodium, phosphate, total bilirubin and total protein.

## Data availability

All data are available in this manuscript and associated supplementary data.

## Code availability

No specific code was produced within this manuscript.

## Acknowledgements

We thank Sven Reiche (Department of Experimental Animal Facilities and Biorisk Management, Friedrich-Loeffler-Institut, Germany) for providing the TULV 1 monoclonal antibody. We acknowledge Jolanda Kreeft – Voermans, Judith Guldemeester, and Richard Molekamp for the implementation of the panHCPS PCR for ANDV detection. The authors are grateful for the outstanding work of Frazer Bell, Lynn Marion Stevenson, Lynn Oxford, Lewis Kidd and Leah Howie from Histology Research Service at the MRC-University of Glasgow, Scotland. We would also like to thank Frans Meijers for his support and advice on project licenses.

## Author contributions

MR (Conceptualization, Project Administration, Data Curation, Formal Analysis, Investigation, Methodology, Visualization, Writing - Original Draft); MW (Data Curation, Formal Analysis, Investigation, Methodology, Writing – Review & Editing); VH (Conceptualization, Data Curation, Formal Analysis, Investigation, Methodology, Supervision, Visualization, Writing - Review & Editing); KH (Data Curation, Formal Analysis, Investigation, Methodology, Writing – Review & Editing); FF (Data Curation, Formal Analysis, Investigation, Methodology); GI (Data Curation, Formal Analysis, Investigation, Methodology); W-WJ (Data Curation, Formal Analysis, Investigation, Methodology); LZ (Data Curation, Formal Analysis, Investigation, Methodology); DvES (Investigation, Methodology); VV (Investigation, Methodology); SS (Investigation, Methodology); JR (Investigation, Methodology); LvD (Investigation, Methodology); MarP (Investigation, Methodology); MtM (Investigation, Methodology); DdM (Investigation, Methodology); HC (Investigation, Methodology, Data Curation, Formal Analysis); KW (Investigation, Methodology); SS-R (Investigation, Methodology); AG (Investigation, Resources); AS (Investigation, Methodology, Supervision, Funding); BOM (Investigation, Supervision); CR (Investigation, Resources, Funding Supervision, Writing – Review & Editing); BLH (Conceptualization, Project Administration, Funding acquisition, Supervision, Writing – review & editing); RdV (Conceptualization, Project Administration, Formal analysis, Funding acquisition, Supervision, Visualization, Writing – review & editing), SH (Conceptualization, Project administration, Funding acquisition, Supervision, Writing – review & editing), MP (Conceptualization, Project administration, Funding acquisition, Supervision, Writing – original draft). MW, VH, KH, FF made an equal contribution to this study. BLH, RdV, SH, MP are joint senior authors.

## Supplementary material

**Supplemental figure S1.**
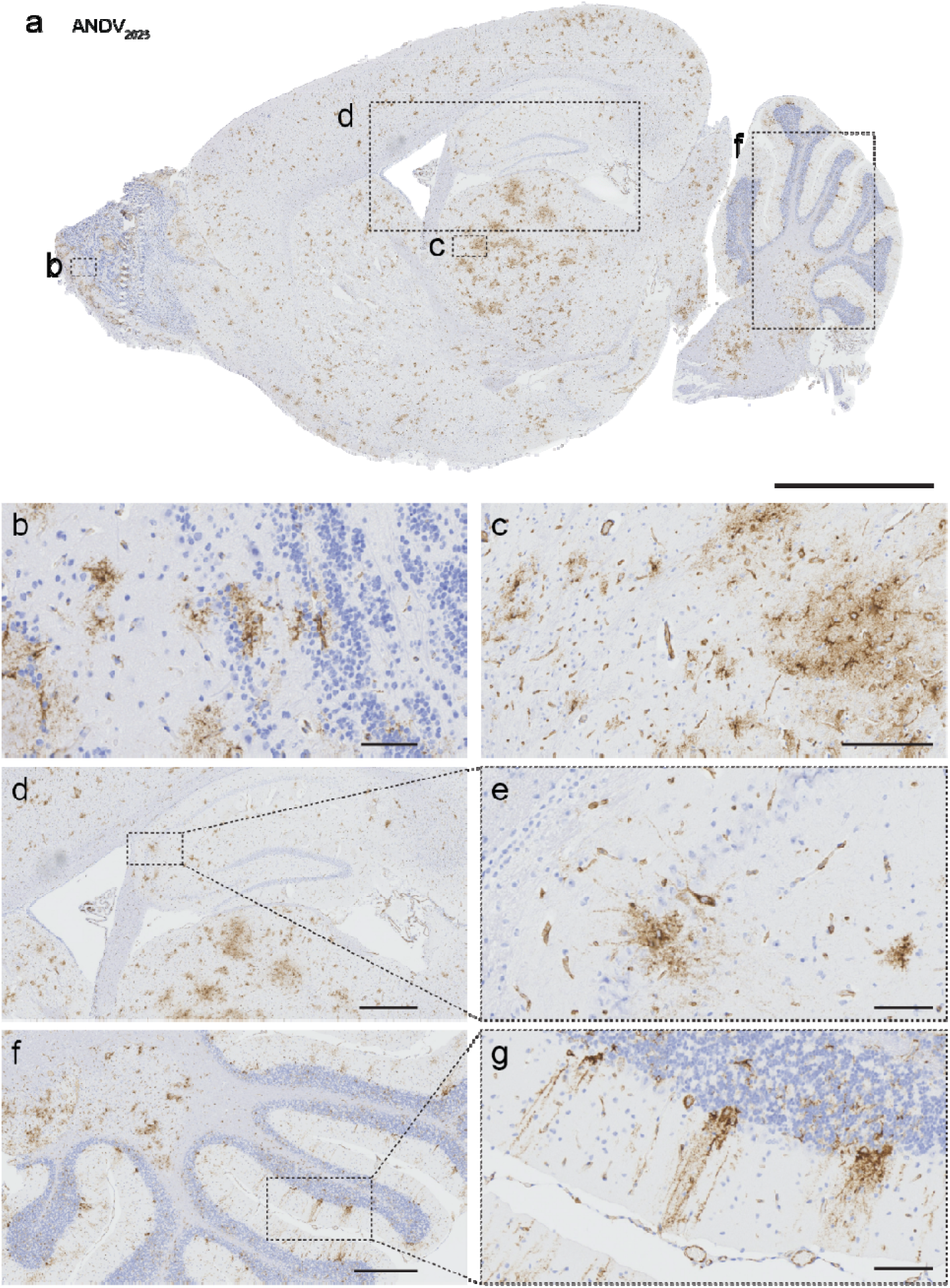
ANDV_2026_ in the brain at 14 dpi. (**a**) Widespread distribution of viral N protein of an ANDV_2026_ inoculated hamster in all areas of the brain (bar, 3 mm). (**b**) Positive cells in the olfactory bulb (bar, 60 µm). (**c**) Positive cells in the mesencephalon (bar, 200 µm). (**d**) Low magnification of the hippocampus (bar, 700 µm). (e) High magnification of positive neurons and endothelial cells in the hippocampus (bar, 80 µm) (**f**) Low magnification of the cerebellum (bar, 500 µm). (**g**) High magnification of virus-positive Purkinje cells in the cerebellum (bar, 90 µm).

**Supplemental figure 2.**
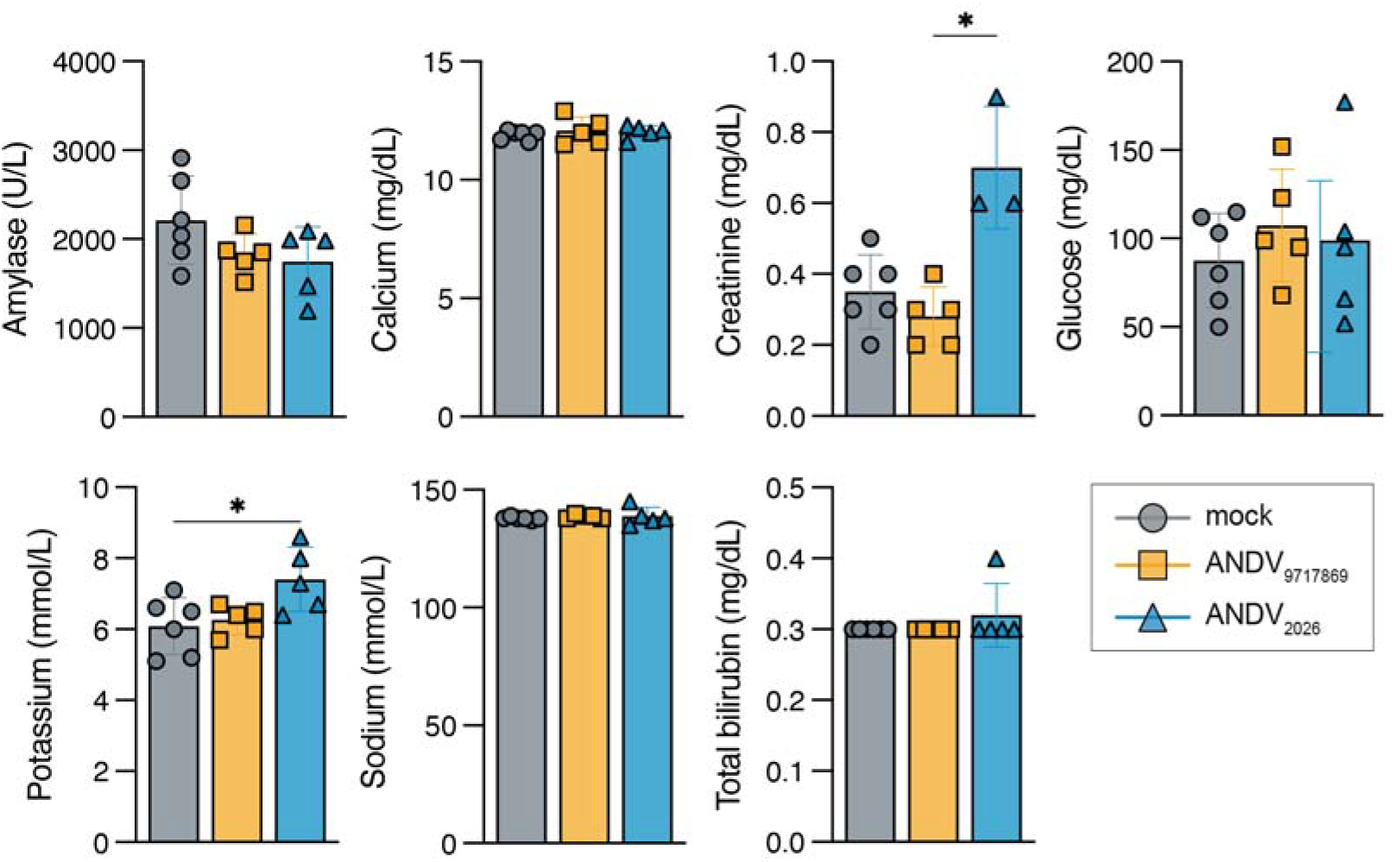
Clinical chemistry of ANDV inoculated hamsters on 12-14 dpi. Data shown as mean ± SD; symbols represent individual hamsters. Data was compared by one-way ANOVA with multiple comparisons; * p<0.05, ** p<0.01, *** p<0.001, **** p<0.0001.

**Supplemental figure 3.**
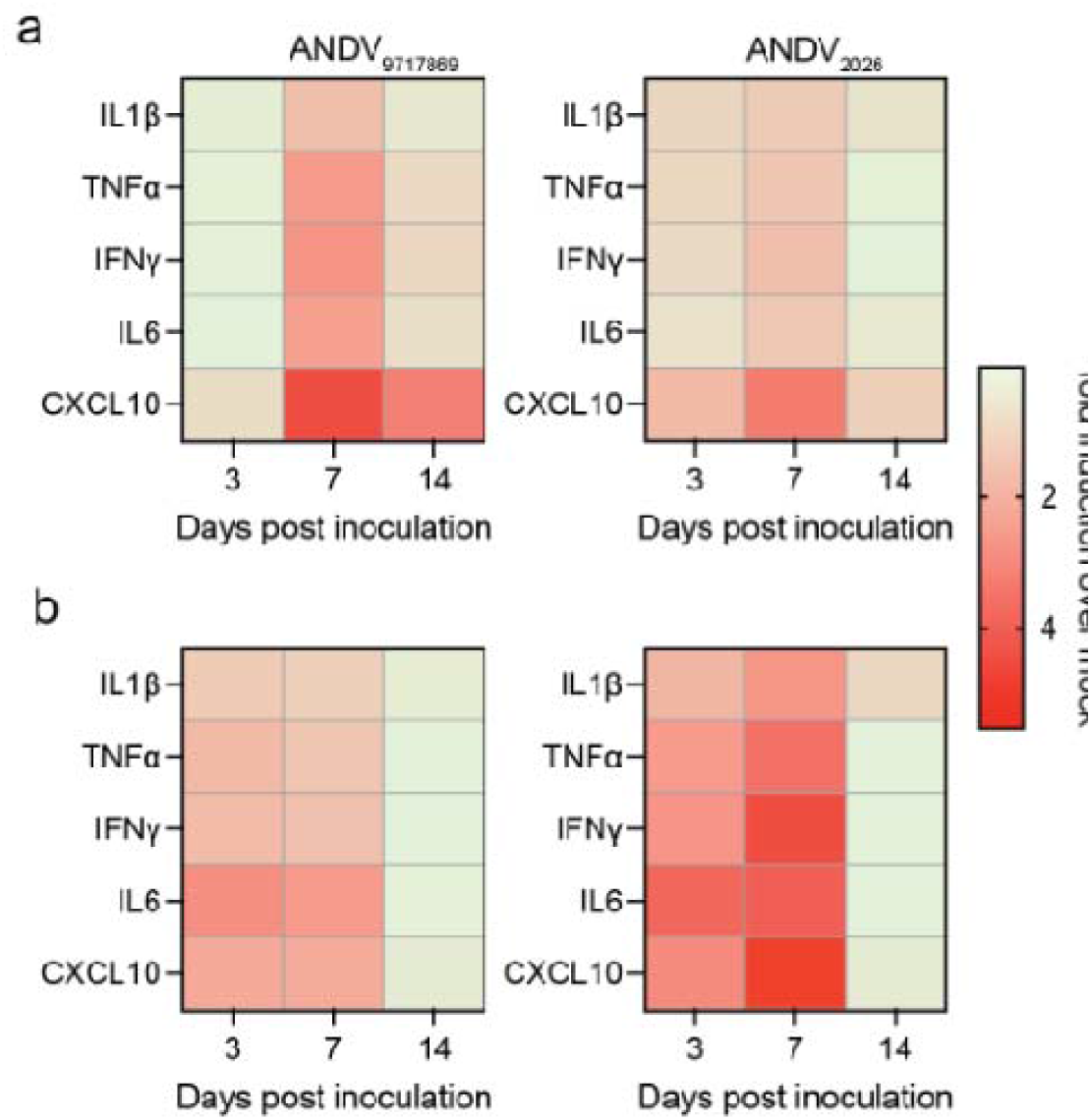
Hostgene response of golden Syrian hamsters after ANDV inoculation. (**a**) Interleukin-1-beta (IL1β), Tumor necrosis factor-alpha (TNFα), interferon gamma (IFNγ), interleukin-6 (IL6) and C-X-C motif chemokine ligand 10 (CXCL-10) were measured via qRT-PCR in lungs on 3, 7 and 12-14 dpi of ANDV inoculated hamsters and were calculated relative to values of mock inoculated animals. (**b**) Interleukin-1-beta (IL1β), Tumor necrosis factor-alpha (TNFα), interferon gamma (IFNγ), interleukin-6 (IL6) and C-X-C motif chemokine ligand 10 (CXCL-10) were measured via qRT-PCR in spleens on 3, 7 and 12-14 dpi of ANDV inoculated hamsters and were calculated relative to values of mock inoculated animals.

**Supplemental figure 4.**
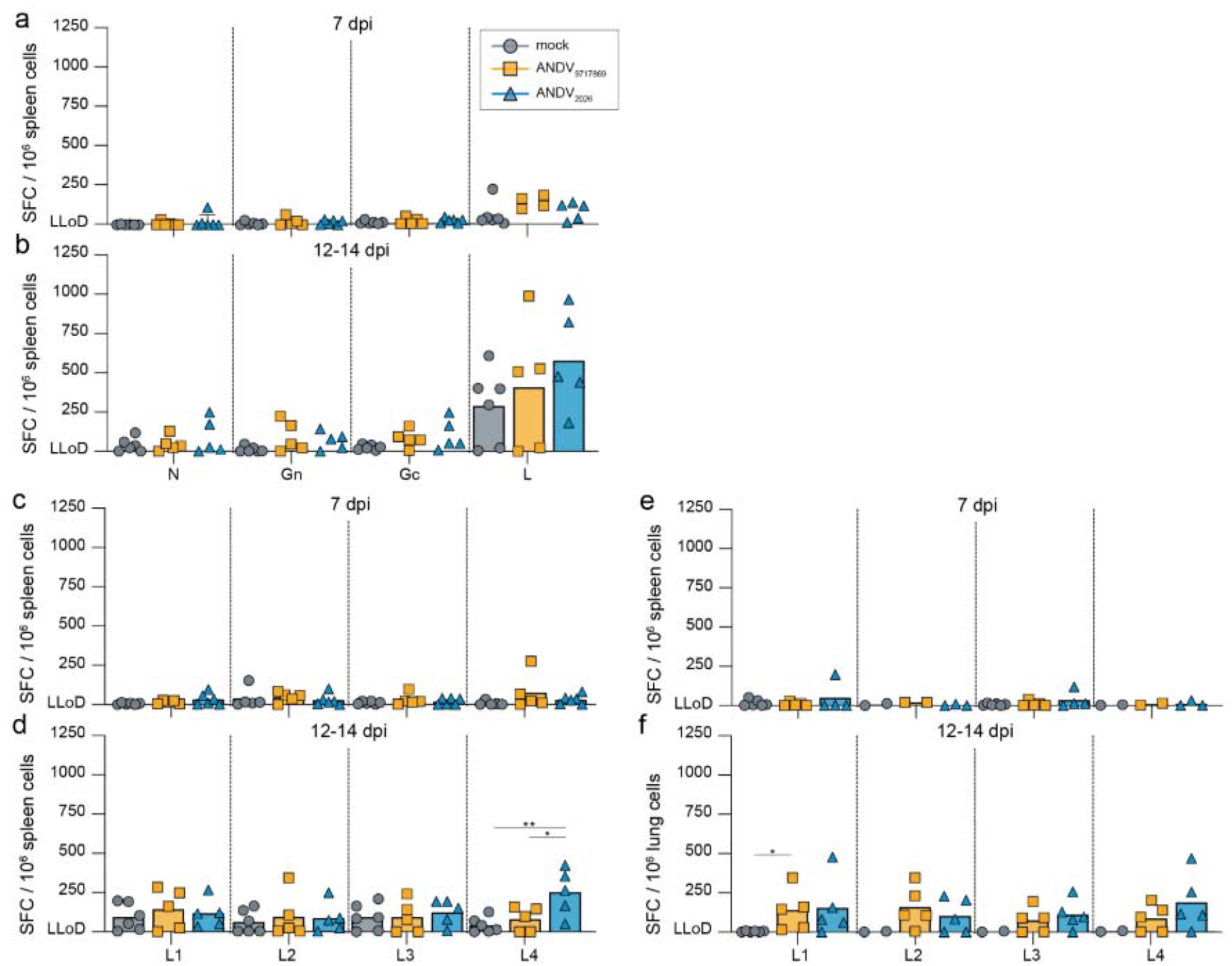
ANDV-specific T cell responses in experimentally inoculated hamsters. (**a,b**) ANDV-specific T cell responses in the spleens measured by stimulation of lymphocytes with overlapping peptide pools and IFNg ELISPOT at 7 and 12-14 dpi. T cell responses are expressed as spot forming cells (SFC) per 10^6^ total cells. T cell responses targeting L are shown as the sum of L1-, L2-, L3-, and L4-specific responses. (**c,d**) T cell responses targeting L are shown separately for L1-, L2-, L3-, and L4-specific responses in spleens. (**e,f**) T cell responses targeting L are shown separately for L1-, L2-, L3-, and L4-specific responses in the lung. (**a-f**) Data shown as mean ± SD; symbols represent individual hamsters. Log-transformed ANDV_9717869_ and ANDV_2026_ data were compared to mock by Kruskal-Wallis or one-way ANOVA with multiple comparisons for non-normal or normal distributed data, respectively; * p<0.05, ** p<0.01. Abbreviations: lower limit of detection (LLoD); virus neutralization (VN); nucleoprotein (N); glycoprotein N (Gn); glycoprotein C (Gc); polymerase (L); days post inoculation (dpi).

**Supplemental figure 5.**
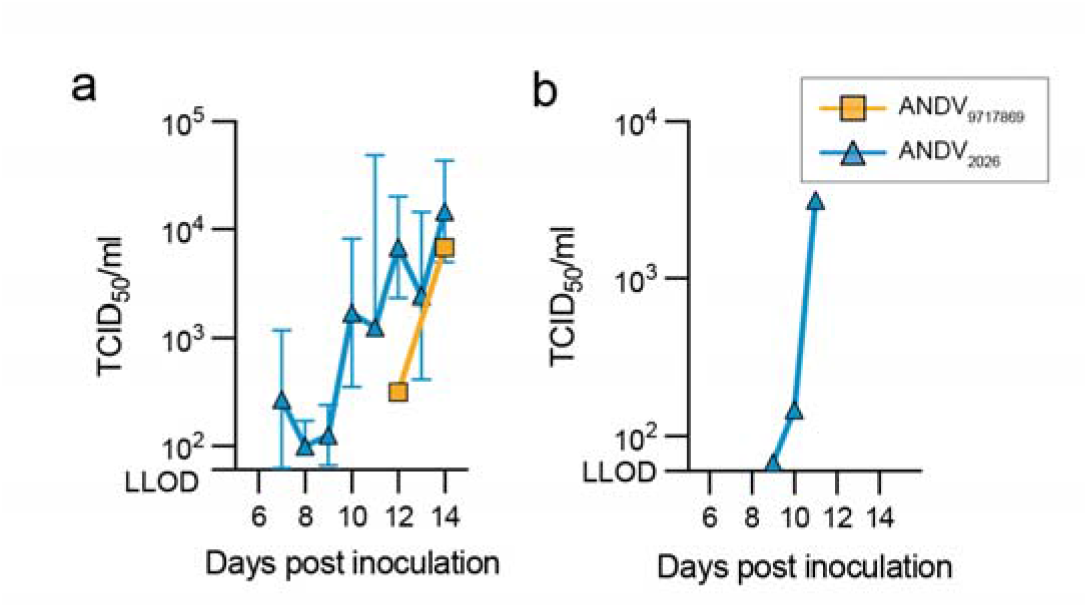
Infectious virus titers in throat swab and feces of ANDV inoculated hamsters. (**a**) Quantification of viral load tested in throat swabs measured by TCID_50_ titration on VeroE6 cells. Only qRT-PCR positive samples (as shown in Fig 6) were subjected to virus titrations. Data shown as geometric mean ± geometric SD; symbols represent individual hamsters. (**b**) Quantification of viral load tested in feces homogenates measured by TCID_50_ titration on VeroE6 cells. Only qRT-PCR positive samples were subjected to virus titrations. Data shown as geometric mean ± geometric SD; symbols represent one cage.

**Supplemental Figure 6:**
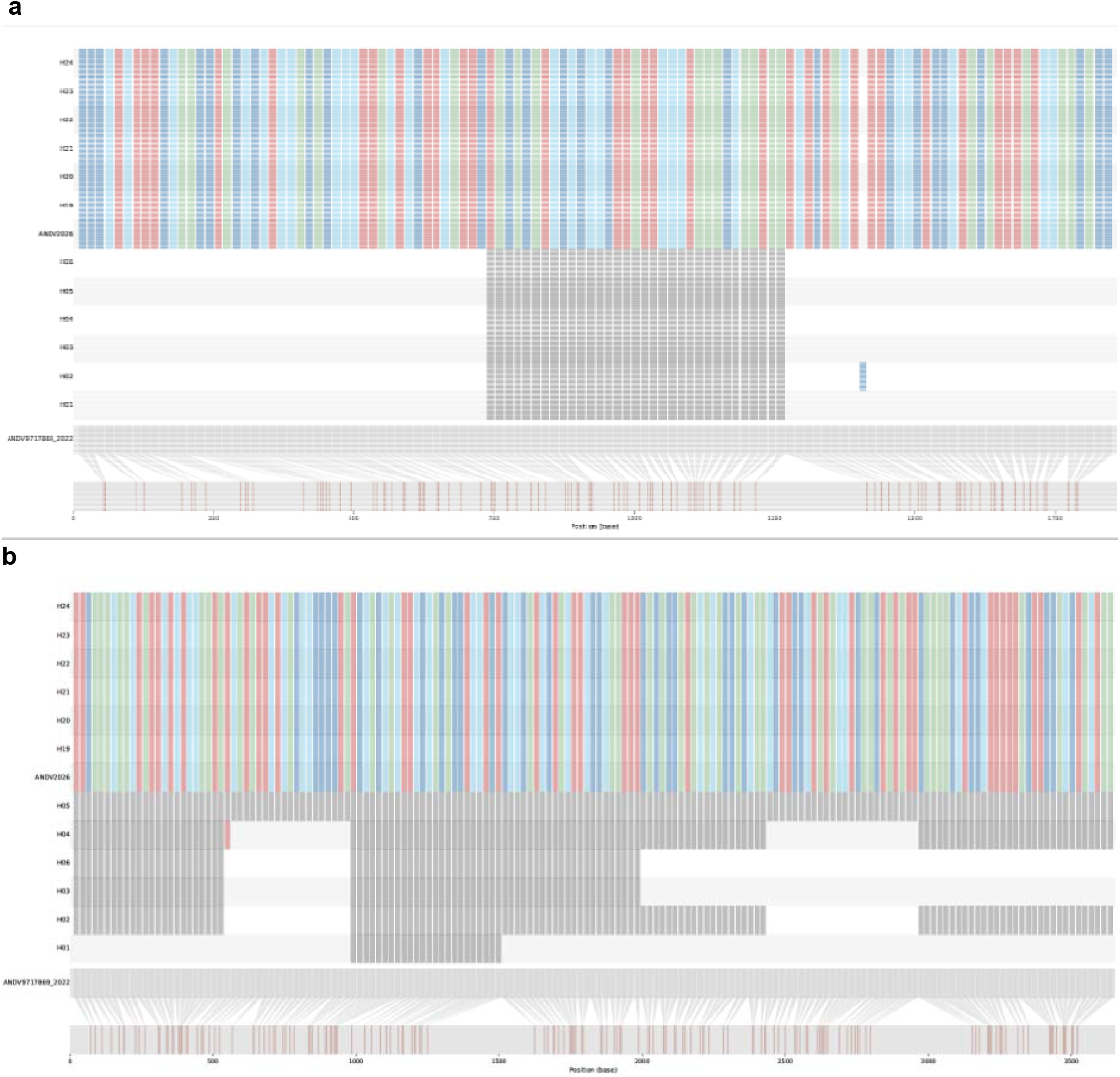
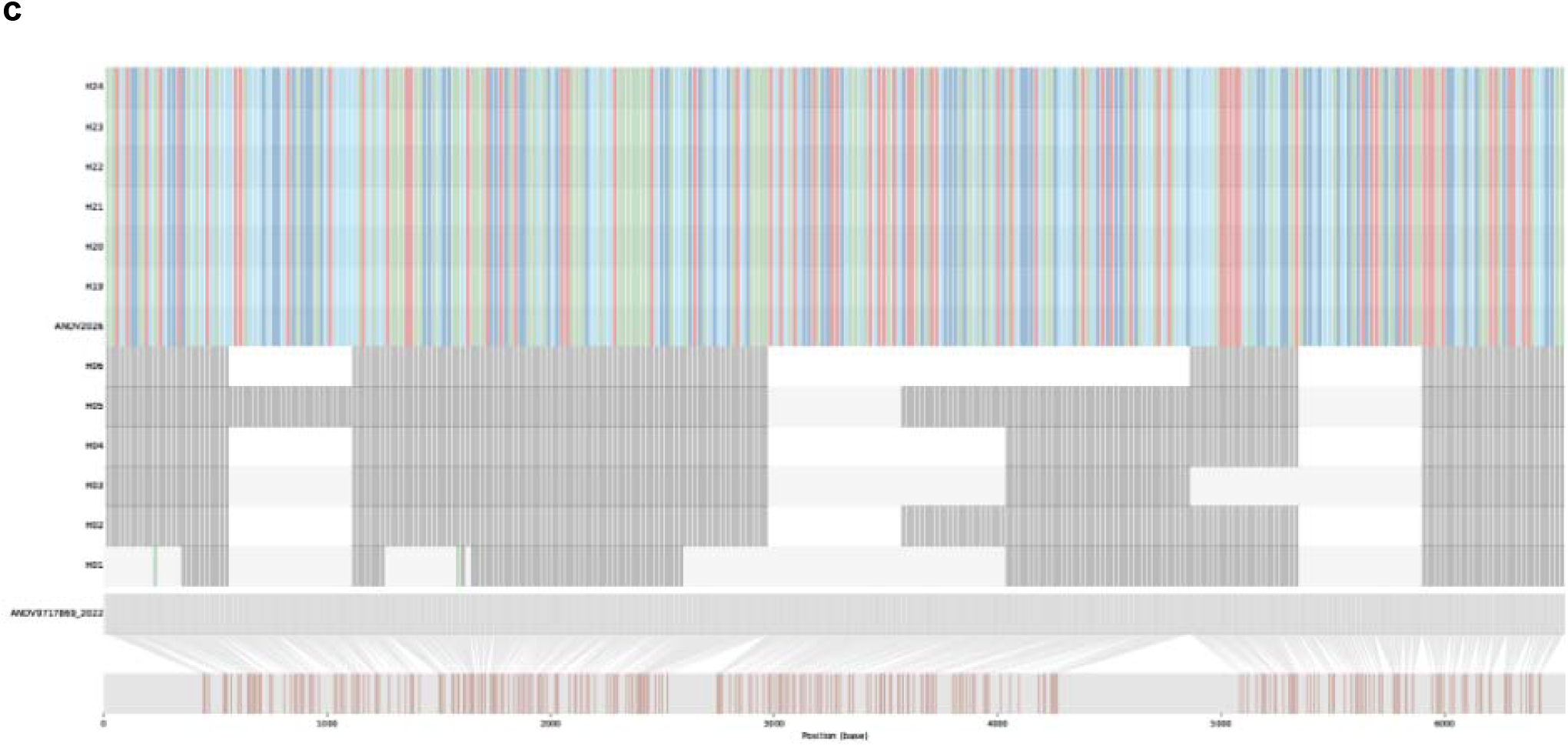
Comparison of single-nucleotide polymorphisms (SNP). Comparison of single-nucleotide polymorphism (SNP) differences within the (**a**) L segment, (**b)** M segment and (**c**) S segment from hamster nasal turbinates (H1-H6, H19-H24), along with control strains from 2020, 2022 and 2026 using amplicon sequencing. SNP changes are shown in comparison to the earliest sequence (ANDV-9717869_2020_V1). Minimum coverage was set to x20 reads. The different colours are representative of the different nucleotide bases: green is T, red is C, dark blue is A, light blue is G and N is dark grey. N indicates positions with a coverage less than 20 reads. White or light grey background indicates a nucleotide identical to the reference sequence (ANDV-9717869_2020_V1).

