## Supplementary tables for "Andes virus from the *MV Hondius* outbreak is highly pathogenic in hamsters and airborne shed"

**Supplementary table 1. Summary of sequencing results.**

| Sample number | Sample type | ANDV virus detected | Ct value | % coverage (x20) L segment | % coverage (x20) M segment | % coverage (x20) S segment |
|---|---|---|---|---|---|---|
| H01 | homogenized tissue material (nasal turbinates) | ANDV $_{9717869}$ | 24.7 | 35.8 | 66.27 | 64.68 |
| H02 | homogenized tissue material (nasal turbinates) | ANDV $_{9717869}$ | 28.43 | 18.98 | 22.39 | 64.68 |
| H03 | homogenized tissue material (nasal turbinates) | ANDV $_{9717869}$ | 24.44 | 30.94 | 44.61 | 64.78 |
| H04 | homogenized tissue material (nasal turbinates) | ANDV $_{9717869}$ | 25.38 | 24.89 | 22.39 | 64.78 |
| H05 | homogenized tissue material (nasal turbinates) | ANDV $_{9717869}$ | 24.95 | 12.64 | 0 | 64.78 |
| H06 | homogenized tissue material (nasal turbinates) | ANDV $_{9717869}$ | 28.8 | 36.42 | 44.61 | 64.62 |
| H19 | homogenized tissue material (nasal turbinates) | ANDV $_{2026}$ | 24.69 | 98.24 | 95.54 | 94.41 |
| H20 | homogenized tissue material (nasal turbinates) | ANDV $_{2026}$ | 26.53 | 98.24 | 95.54 | 94.41 |
| H21 | homogenized tissue material (nasal turbinates) | ANDV $_{2026}$ | 27.41 | 98.24 | 95.54 | 94.41 |
| H22 | homogenized tissue material (nasal turbinates) | ANDV $_{2026}$ | 25.91 | 98.24 | 95.54 | 94.41 |
| H23 | homogenized tissue material (nasal turbinates) | ANDV $_{2026}$ | 27.21 | 98.24 | 95.54 | 94.41 |
| H24 | homogenized tissue material (nasal turbinates) | ANDV $_{2026}$ | 27.87 | 98.24 | 95.54 | 94.41 |
| ANDV-9717869_2022_V2 | virus stocks | ANDV | 15-17 | 77.43 | 76.9 | 86.24 |
| ANDV-2026_V3 | virus stocks | ANDV | 15-17 | 98.24 | 95.54 | 94.41 |

### Supplementary table 2. Peptide pools.

| | | | | | | | | | | | | | |
|---|---|---|---|---|---|---|---|---|---|---|---|---|---|
| **ANDV-N** | MSTLQELQENITAHE | **ANDV-Gc** | MEGWYLVALGVCYTL | **ANDV-Gn** | AETPLMESGWSDTAH | **ANDV-L1** | MEKYREIHQRVRDLA | **ANDV-L2** | IGLIDADNIDSKAEI | **ANDV-L3** | AAVSLLFREVWKQLF | **ANDV-L4** | KEVLKAMSLYSVKEN |
| **ANDV-N** | ELQENITAHEQQLVT | **ANDV-Gc** | LVALGVCYTLTLAMP | **ANDV-Gn** | MESGWSDTAHGVGEI | **ANDV-L1** | EIHQRVRDLAPGTVS | **ANDV-L2** | ADNIDSKAEIDGVTW | **ANDV-L3** | LFREVWKQLFPELEC | **ANDV-L4** | AMSLYSVKENIGLQD |
| **ANDV-N** | ITAHEQQLVTARQKL | **ANDV-Gc** | VCYTLTLAMPKTIYE | **ANDV-Gn** | SDTAHGVGEIPMKTD | **ANDV-L1** | VRDLAPGTVSALECI | **ANDV-L2** | SKAEIDGVTWCYSKV | **ANDV-L3** | WKQLFPELECFFEFA | **ANDV-L4** | SVKENIGLQDDIAAT |
| **ANDV-N** | QQLVTARQKLKDAEK | **ANDV-Gc** | TLAMPKTIYELKMEC | **ANDV-Gn** | GVGEIPMKTDLELDF | **ANDV-L1** | PGTVSALECIDLLDR | **ANDV-L2** | DGVTWCYSKVMSIDL | **ANDV-L3** | PELECFFEFAHHSDD | **ANDV-L4** | IGLQDDIAATRLCIE |
| **ANDV-N** | ARQKLKDAEKAVEVD | **ANDV-Gc** | KTIYELKMECPHTVG | **ANDV-Gn** | PMKTDLELDFSLPSS | **ANDV-L1** | ALECIDLLDRLYAVR | **ANDV-L2** | CYSKVMSIDLNRLLA | **ANDV-L3** | FFEFAHHSDDALFIY | **ANDV-L4** | DIAATRLCIEIWRWC |
| **ANDV-N** | KDAEKAVEVDPDDVN | **ANDV-Gc** | LKMECPHTVGLGQGY | **ANDV-Gn** | LELDFSLPSSSSYSY | **ANDV-L1** | DLLDRLYAVRHDLVD | **ANDV-L2** | MSIDLNRLLALNIAF | **ANDV-L3** | HHSDDALFIYGYLEP | **ANDV-L4** | RLCIEIWRWCKANEQ |
| **ANDV-N** | AVEVDPDDVNKSTLQ | **ANDV-Gc** | PHTVGLGQGYIIGST | **ANDV-Gn** | SLPSSSSYSYRRKLT | **ANDV-L1** | LYAVRHDLVDQMIKH | **ANDV-L2** | NRLLALNIAFEKSLL | **ANDV-L3** | ALFIYGYLEPEDDGT | **ANDV-L4** | IWRWCKANEQDVKEW |
| **ANDV-N** | PDDVNKSTLQSRRAA | **ANDV-Gc** | LGQGYIIGSTELGLI | **ANDV-Gn** | SSYSYRRKLTNPANK | **ANDV-L1** | HDLVDQMIKHDWSDN | **ANDV-L2** | LNIAFEKSLLATATW | **ANDV-L3** | GYLEPEDDGTDWFLY | **ANDV-L4** | KANEQDVKEWLTSLY |
| **ANDV-N** | KSTLQSRRAAVSTLE | **ANDV-Gc** | IIGSTELGLISIEAA | **ANDV-Gn** | RRKLTNPANKEESIP | **ANDV-L1** | QMIKHDWSDNKDVER | **ANDV-L2** | EKSLLATATWFQYYT | **ANDV-L3** | EDDGTDWFLYVSQQI | **ANDV-L4** | DVKEWLTSLYFEKQT |
| **ANDV-N** | SRRAAVSTLETKLGE | **ANDV-Gc** | ELGLISIEAASDIKL | **ANDV-Gn** | NPANKEESIPFHFQM | **ANDV-L1** | DWSDNKDVERPIGQV | **ANDV-L2** | ATATWFQYYTEDQGH | **ANDV-L3** | DWFLYVSQQIQAGNF | **ANDV-L4** | LTSLYFEKQTLMDWV |
| **ANDV-N** | VSTLETKLGELKRQL | **ANDV-Gc** | SIEAASDIKLESSCN | **ANDV-Gn** | EESIPFHFQMEKQVI | **ANDV-L1** | KDVERPIGQVLLMAG | **ANDV-L2** | FQYYTEDQGHFPLQH | **ANDV-L3** | VSQQIQAGNFHWHAI | **ANDV-L4** | FEKQTLMDWVERFRR |
| **ANDV-N** | TKLGELKRQLADLVA | **ANDV-Gc** | SDIKLESSCNFDLHT | **ANDV-Gn** | FHFQMEKQVIHAEIQ | **ANDV-L1** | PIGQVLLMAGIPNDI | **ANDV-L2** | EDQGHFPLQHALRSI | **ANDV-L3** | QAGNFHWHAINQEMW | **ANDV-L4** | LMDWVERFRRKGVVP |
| **ANDV-N** | LKRQLADLVAAQKLA | **ANDV-Gc** | ESSCNFDLHTTSMAQ | **ANDV-Gn** | EKQVIHAEIQPLGHW | **ANDV-L1** | LLMAGIPNDIIQGME | **ANDV-L2** | FPLQHALRSIFSFHF | **ANDV-L3** | HWHAINQEMWKSMFN | **ANDV-L4** | ERFRRKGVVPIDPEI |
| **ANDV-N** | ADLVAAQKLATKPVD | **ANDV-Gc** | FDLHTTSMAQKSFTQ | **ANDV-Gn** | HAEIQPLGHWMDATF | **ANDV-L1** | IPNDIIQGMEKKIIP | **ANDV-L2** | ALRSIFSFHFLLCVS | **ANDV-L3** | NQEMWKSMFNLHEHL | **ANDV-L4** | KGVVPIDPEIQCIGL |
| **ANDV-N** | AQKLATKPVDPTGLE | **ANDV-Gc** | TSMAQKSFTQVEWRK | **ANDV-Gn** | PLGHWMDATFNIKTA | **ANDV-L1** | IQGMEKKIIPNSPSG | **ANDV-L2** | FSFHFLLCVSQKMKL | **ANDV-L3** | KSMFNLHEHLLLMGS | **ANDV-L4** | IDPEIQCIGLLLYDV |
| **ANDV-N** | TKPVDPTGLEPDDHL | **ANDV-Gc** | KSFTQVEWRKKSDTT | **ANDV-Gn** | MDATFNIKTAFHCYG | **ANDV-L1** | KKIIPNSPSGQVLKS | **ANDV-L2** | LLCVSQKMKLCAIFD | **ANDV-L3** | LHEHLLLMGSIKVSP | **ANDV-L4** | QCIGLLLYDVLGYKS |
| **ANDV-N** | PTGLEPDDHLKEKSS | **ANDV-Gc** | VEWRKKSDTTDTTNA | **ANDV-Gn** | NIKTAFHCYGACQKY | **ANDV-L1** | NSPSGQVLKSFFRMT | **ANDV-L2** | QKMKLCAIFDNLRYL | **ANDV-L3** | LLMGSIKVSPKKTTV | **ANDV-L4** | LLYDVLGYKSVLQMQ |
| **ANDV-N** | PDDHLKEKSSLRYGN | **ANDV-Gc** | KSDTTDTTNAASTTF | **ANDV-Gn** | FHCYGACQKYSYPWQ | **ANDV-L1** | QVLKSFFRMTPDNYK | **ANDV-L2** | CAIFDNLRYLIPSVT | **ANDV-L3** | IKVSPKKTTVSPTNA | **ANDV-L4** | LGYKSVLQMQANRRA |
| **ANDV-N** | KEKSSLRYGNVLDVN | **ANDV-Gc** | DTTNAASTTFEAQTK | **ANDV-Gn** | ACQKYSYPWQTSKCF | **ANDV-L1** | FFRMTPDNYKITGNL | **ANDV-L2** | NLRYLIPSVTSLYSG | **ANDV-L3** | KKTTVSPTNAEFLST | **ANDV-L4** | VLQMQANRRAYSGKQ |
| **ANDV-N** | LRYGNVLDVNSIDLE | **ANDV-Gc** | ASTTFEAQTKTVNLR | **ANDV-Gn** | SYPWQTSKCFFEKDY | **ANDV-L1** | PDNYKITGNLIEFIE | **ANDV-L2** | IPSVTSLYSGYELLI | **ANDV-L3** | SPTNAEFLSTFFEGC | **ANDV-L4** | ANRRAYSGKQYDAYC |
| **ANDV-N** | VLDVNSIDLEEPSGQ | **ANDV-Gc** | EAQTKTVNLRGTCVL | **ANDV-Gn** | TSKCFFEKDYQYETG | **ANDV-L1** | ITGNLIEFIEVTVTA | **ANDV-L2** | SLYSGYELLIEKFFE | **ANDV-L3** | EFLSTFFEGCAVSIP | **ANDV-L4** | YSGKQYDAYCVQTYN |
| **ANDV-N** | SIDLEEPSGQTADWK | **ANDV-Gc** | TVNLRGTCVLAPELY | **ANDV-Gn** | FEKDYQYETGWGCNP | **ANDV-L1** | IEFIEVTVTADVSRG | **ANDV-L2** | YELLIEKFFERPFKS | **ANDV-L3** | FFEGCAVSIPFVKIL | **ANDV-L4** | YDAYCVQTYNEETKL |
| **ANDV-N** | EPSGQTADWKAIGAY | **ANDV-Gc** | GTCVLAPELYDTLKK | **ANDV-Gn** | QYETGWGCNPGDCPG | **ANDV-L1** | VTVTADVSRGIREKK | **ANDV-L2** | EKFFERPFKSSLDVY | **ANDV-L3** | AVSIPFVKILLGSLS | **ANDV-L4** | VQTYNEETKLYEGDL |
| **ANDV-N** | TADWKAIGAYILGFA | **ANDV-Gc** | APELYDTLKKVKKTV | **ANDV-Gn** | WGCNPGDCPGVGTGC | **ANDV-L1** | DVSRGIREKKIKYEG | **ANDV-L2** | RPFKSSLDVYLYSII | **ANDV-L3** | FVKILLGSLSDLPGL | **ANDV-L4** | EETKLYEGDLRVTFN |
| **ANDV-N** | AIGAYILGFAIPIIL | **ANDV-Gc** | DTLKKVKKTVLCYDL | **ANDV-Gn** | GDCPGVGTGCTACGV | **ANDV-L1** | IREKKIKYEGGLQFV | **ANDV-L2** | SLDVYLYSIIKSLLI | **ANDV-L3** | LGSLSDLPGLGFFDD | **ANDV-L4** | YEGDLRVTFNFGLDC |
| **ANDV-N** | ILGFAIPIILKALYM | **ANDV-Gc** | VKKTVLCYDLTCNQT | **ANDV-Gn** | VGTGCTACGVYLDKL | **ANDV-L1** | IKYEGGLQFVEHLLE | **ANDV-L2** | LYSIIKSLLISLAQN | **ANDV-L3** | DLPGLGFFDDLAAAQ | **ANDV-L4** | RVTFNFGLDCARLEV |
| **ANDV-N** | IPIILKALYMLSTRG | **ANDV-Gc** | LCYDLTCNQTHCQPT | **ANDV-Gn** | TACGVYLDKLKSVGK | **ANDV-L1** | GLQFVEHLLETESRK | **ANDV-L2** | KSLLISLAQNNKVRF | **ANDV-L3** | GFFDDLAAAQSRCVK | **ANDV-L4** | FGLDCARLEVFWDKK |

| | | | | | | | | | | | | | |
|---|---|---|---|---|---|---|---|---|---|---|---|---|---|
| **ANDV-N** | KALYMLSTRGRQTVK | **ANDV-Gc** | TCNQTHCQPTVYLIA | **ANDV-Gn** | YLDKLKSVGKAYKII | **ANDV-L1** | EHLLETESRKGNIPQ | **ANDV-L2** | SLAQNNKVRFYSRVR | **ANDV-L3** | LAAAQSRCVKSLDLG | **ANDV-L4** | ARLEVFWDKKEYILE |
| **ANDV-N** | LSTRGRQTVKDNKGT | **ANDV-Gc** | HCQPTVYLIAPVLTC | **ANDV-Gn** | KSVGKAYKIISLKYT | **ANDV-L1** | TESRKGNIPQPYKIT | **ANDV-L2** | NKVRFYSRVRLLGLT | **ANDV-L3** | SRCVKSLDLGACPQL | **ANDV-L4** | FWDKKEYILETSITQ |
| **ANDV-N** | RQTVKDNKGTRIRFK | **ANDV-Gc** | VYLIAPVLTCMSIRS | **ANDV-Gn** | AYKIISLKYTRKVCI | **ANDV-L1** | GNIPQPYKITFSVVA | **ANDV-L2** | YSRVRLLGLTVDHST | **ANDV-L3** | SLDLGACPQLAQLAI | **ANDV-L4** | EYILETSITQRHVLR |
| **ANDV-N** | DNKGTRIRFKDDSSF | **ANDV-Gc** | PVLTCMSIRSCMASV | **ANDV-Gn** | SLKYTRKVCIQLGTE | **ANDV-L1** | PYKITFSVVAVKTDG | **ANDV-L2** | LLGLTVDHSTVGASG | **ANDV-L3** | ACPQLAQLAIVLCTS | **ANDV-L4** | TSITQRHVLRLLMEE |
| **ANDV-N** | RIRFKDDSSFEEVNG | **ANDV-Gc** | MSIRSCMASVFTSRI | **ANDV-Gn** | RKVCIQLGTEQTCKH | **ANDV-L1** | FSVVAVKTDGSNIST | **ANDV-L2** | VDHSTVGASGVYPSL | **ANDV-L3** | AQLAIVLCTSKVERL | **ANDV-L4** | RHVLRLLMEEVSQEL |
| **ANDV-N** | DDSSFEEVNGIRKPK | **ANDV-Gc** | CMASVFTSRIQVIYE | **ANDV-Gn** | QLGTEQTCKHIDAND | **ANDV-L1** | VKTDGSNISTQWPSR | **ANDV-L2** | VGASGVYPSLMSRVV | **ANDV-L3** | VLCTSKVERLYGTAD | **ANDV-L4** | LLMEEVSQELIRCGM |
| **ANDV-N** | EEVNGIRKPKHLYVS | **ANDV-Gc** | FTSRIQVIYEKTHCV | **ANDV-Gn** | QTCKHIDANDCLVTP | **ANDV-L1** | SNISTQWPSRRNDGV | **ANDV-L2** | VYPSLMSRVVYKHYR | **ANDV-L3** | KVERLYGTADGMVNS | **ANDV-L4** | VSQELIRCGMRFKTE |
| **ANDV-N** | IRKPKHLYVSMPTAQ | **ANDV-Gc** | QVIYEKTHCVTGQLI | **ANDV-Gn** | IDANDCLVTPSVKVC | **ANDV-L1** | QWPSRRNDGVVQHMR | **ANDV-L2** | MSRVVYKHYRSLISE | **ANDV-L3** | YGTADGMVNSPTAFL | **ANDV-L4** | IRCGMRFKTEQVNQT |
| **ANDV-N** | HLYVSMPTAQSTMKA | **ANDV-Gc** | KTHCVTGQLIEGQCF | **ANDV-Gn** | CLVTPSVKVCIVGTV | **ANDV-L1** | RNDGVVQHMRLVQAD | **ANDV-L2** | YKHYRSLISEATTCF | **ANDV-L3** | GMVNSPTAFLKVNKA | **ANDV-L4** | RFKTEQVNQTRSLVL |
| **ANDV-N** | MPTAQSTMKAEEITP | **ANDV-Gc** | TGQLIEGQCFNPAHT | **ANDV-Gn** | SVKVCIVGTVSKLQP | **ANDV-L1** | VQHMRLVQADINYVR | **ANDV-L2** | SLISEATTCFFLFEK | **ANDV-L3** | PTAFLKVNKAHVPVP | **ANDV-L4** | QVNQTRSLVLFKTEA |
| **ANDV-N** | STMKAEEITPGRFRT | **ANDV-Gc** | EGQCFNPAHTLTLSQ | **ANDV-Gn** | IVGTVSKLQPSDTLL | **ANDV-L1** | LVQADINYVREHLIK | **ANDV-L2** | ATTCFFLFEKGLHGN | **ANDV-L3** | KVNKAHVPVPLGGDG | **ANDV-L4** | RSLVLFKTEAGFEWG |
| **ANDV-N** | EEITPGRFRTIACGL | **ANDV-Gc** | NPAHTLTLSQPAHTY | **ANDV-Gn** | SKLQPSDTLLFLGPL | **ANDV-L1** | INYVREHLIKLDERA | **ANDV-L2** | FLFEKGLHGNLPEEA | **ANDV-L3** | HVPVPLGGDGSMSIM | **ANDV-L4** | FKTEAGFEWGKPNVP |
| **ANDV-N** | GRFRTIACGLFPAQV | **ANDV-Gc** | LTLSQPAHTYDTVTL | **ANDV-Gn** | SDTLLFLGPLEQGGI | **ANDV-L1** | EHLIKLDERASLEAM | **ANDV-L2** | GLHGNLPEEAKIHLE | **ANDV-L3** | LGGDGSMSIMELATA | **ANDV-L4** | GFEWGKPNVPCIVYK |
| **ANDV-N** | IACGLFPAQVKARNI | **ANDV-Gc** | PAHTYDTVTLPISCF | **ANDV-Gn** | FLGPLEQGGIILKQW | **ANDV-L1** | LDERASLEAMFNLKF | **ANDV-L2** | LPEEAKIHLETIEWA | **ANDV-L3** | SMSIMELATAGFGMA | **ANDV-L4** | KPNVPCIVYKHCVLR |
| **ANDV-N** | FPAQVKARNIISPVM | **ANDV-Gc** | DTVTLPISCFFTPKK | **ANDV-Gn** | EQGGIILKQWCTTSC | **ANDV-L1** | SLEAMFNLKFHVSGP | **ANDV-L2** | KIHLETIEWARKFQE | **ANDV-L3** | ELATAGFGMADKNIL | **ANDV-L4** | CIVYKHCVLRTGLRT |
| **ANDV-N** | KARNIISPVMGVIGF | **ANDV-Gc** | PISCFFTPKKSEQLK | **ANDV-Gn** | ILKQWCTTSCAFGDP | **ANDV-L1** | FNLKFHVSGPKLRYF | **ANDV-L2** | TIEWARKFQEKEKQY | **ANDV-L3** | GFGMADKNILKNAFI | **ANDV-L4** | HCVLRTGLRTKQPIN |
| **ANDV-N** | ISPVMGVIGFGFFVK | **ANDV-Gc** | FTPKKSEQLKVIKTF | **ANDV-Gn** | CTTSCAFGDPGDIMS | **ANDV-L1** | HVSGPKLRYFNIPDY | **ANDV-L2** | RKFQEKEKQYGDILL | **ANDV-L3** | DKNILKNAFISYKHT | **ANDV-L4** | TGLRTKQPINKEFMI |
| **ANDV-N** | GVIGFGFFVKDWMDR | **ANDV-Gc** | SEQLKVIKTFEGILT | **ANDV-Gn** | AFGDPGDIMSTPSGM | **ANDV-L1** | KLRYFNIPDYRPQQL | **ANDV-L2** | KEKQYGDILLKEGYT | **ANDV-L3** | KNAFISYKHTRRDGD | **ANDV-L4** | KQPINKEFMINVQSD |
| **ANDV-N** | GFFVKDWMDRIEEFL | **ANDV-Gc** | VIKTFEGILTKTGCT | **ANDV-Gn** | GDIMSTPSGMRCPEH | **ANDV-L1** | NIPDYRPQQLCEPRI | **ANDV-L2** | GDILLKEGYTIESVI | **ANDV-L3** | SYKHTRRDGDRYVLG | **ANDV-L4** | KEFMINVQSDGFRAI |
| **ANDV-N** | DWMDRIEEFLAAECP | **ANDV-Gc** | EGILTKTGCTENALQ | **ANDV-Gn** | TPSGMRCPEHTGSFR | **ANDV-L1** | RPQQLCEPRIDNLIQ | **ANDV-L2** | KEGYTIESVINGEVD | **ANDV-L3** | RRDGDRYVLGLFKFL | **ANDV-L4** | NVQSDGFRAIAQMDI |
| **ANDV-N** | IEEFLAAECPFLPKP | **ANDV-Gc** | KTGCTENALQGYYVC | **ANDV-Gn** | RCPEHTGSFRKICGF | **ANDV-L1** | CEPRIDNLIQYCKNW | **ANDV-L2** | IESVINGEVDVEQQL | **ANDV-L3** | RYVLGLFKFLMSLSE | **ANDV-L4** | GFRAIAQMDIESPRF |
| **ANDV-N** | AAECPFLPKPKVASE | **ANDV-Gc** | ENALQGYYVCFLGSH | **ANDV-Gn** | TGSFRKICGFATTPV | **ANDV-L1** | DNLIQYCKNWLTKEH | **ANDV-L2** | NGEVDVEQQLFCQEV | **ANDV-L3** | LFKFLMSLSEDVFQH | **ANDV-L4** | AQMDIESPRFLLAHA |
| **ANDV-N** | FLPKPKVASEAFMST | **ANDV-Gc** | GYYVCFLGSHSEPLI | **ANDV-Gn** | KICGFATTPVCEYQG | **ANDV-L1** | YCKNWLTKEHKFVFK | **ANDV-L2** | VEQQLFCQEVSELSA | **ANDV-L3** | MSLSEDVFQHDRLGE | **ANDV-L4** | ESPRFLLAHAYHTLR |
| **ANDV-N** | KVASEAFMSTNKMYF | **ANDV-Gc** | FLGSHSEPLIVPSLE | **ANDV-Gn** | ATTPVCEYQGNTISG | **ANDV-L1** | LTKEHKFVFKEVSGA | **ANDV-L2** | FCQEVSELSAQELNK | **ANDV-L3** | DVFQHDRLGEFSFVG | **ANDV-L4** | LLAHAYHTLRDIRYQ |
| **ANDV-N** | AFMSTNKMYFLNRQR | **ANDV-Gc** | SEPLIVPSLEDIRSA | **ANDV-Gn** | CEYQGNTISGYKRMM | **ANDV-L1** | KFVFKEVSGANVIQA | **ANDV-L2** | SELSAQELNKYLQAK | **ANDV-L3** | DRLGEFSFVGKVQWK | **ANDV-L4** | YHTLRDIRYQAVQAV |
| **ANDV-N** | NKMYFLNRQRQVNES | **ANDV-Gc** | VPSLEDIRSAEVVSR | **ANDV-Gn** | NTISGYKRMMATKDS | **ANDV-L1** | EVSGANVIQAFESHE | **ANDV-L2** | QELNKYLQAKSQVLC | **ANDV-L3** | FSFVGKVQWKVFTPK | **ANDV-L4** | DIRYQAVQAVGNVWF |
| **ANDV-N** | LNRQRQVNESKVQDI | **ANDV-Gc** | DIRSAEVVSRMLVHP | **ANDV-Gn** | YKRMMATKDSFQSFN | **ANDV-L1** | NVIQAFESHEQLHLQ | **ANDV-L2** | YLQAKSQVLCANIMN | **ANDV-L3** | KVQWKVFTPKAEFEF | **ANDV-L4** | AVQAVGNVWFKTEQH |
| **ANDV-N** | QVNESKVQDIIDLID | **ANDV-Gc** | EVVSRMLVHPRGEDH | **ANDV-Gn** | ATKDSFQSFNLTEPH | **ANDV-L1** | FESHEQLHLQKYNES | **ANDV-L2** | SQVLCANIMNKHWDK | **ANDV-L3** | VFTPKAEFEFHDQFS | **ANDV-L4** | GNVWFKTEQHKLFIN |
| **ANDV-N** | KVQDIIDLIDHAETE | **ANDV-Gc** | MLVHPRGEDHDAIQN | **ANDV-Gn** | FQSFNLTEPHITTNK | **ANDV-L1** | QLHLQKYNESRKPRN | **ANDV-L2** | ANIMNKHWDKPYFSQ | **ANDV-L3** | AEFEFHDQFSHNYLL | **ANDV-L4** | KTEQHKLFINPIISS |

| | | | | | | | | | | | | | |
|---|---|---|---|---|---|---|---|---|---|---|---|---|---|
| **ANDV-N** | IDLIDHAETESATLF | **ANDV-Gc** | RGEDHDAIQNSQSYL | **ANDV-Gn** | LTEPHITTNKLEWID | **ANDV-L1** | KYNESRKPRNFLLLQ | **ANDV-L2** | KHWDKPYFSQTRNIS | **ANDV-L3** | HDQFSHNYLLEWTRQ | **ANDV-L4** | KLFINPIISSGLLEN |
| **ANDV-N** | HAETESATLFTEIAT | **ANDV-Gc** | DAIQNSQSYLRIVGP | **ANDV-Gn** | ITTNKLEWIDPDGNT | **ANDV-L1** | RKPRNFLLLQLTVQG | **ANDV-L2** | PYFSQTRNISLKGMS | **ANDV-L3** | HNYLLEWTRQHPVYD | **ANDV-L4** | PIISSGLLENFMKGL |
| **ANDV-N** | SATLFTEIATPHSVW | **ANDV-Gc** | SQSYLRIVGPITAKV | **ANDV-Gn** | LEWIDPDGNTRDHVN | **ANDV-L1** | FLLLQLTVQGAYLPS | **ANDV-L2** | TRNISLKGMSGALQE | **ANDV-L3** | EWTRQHPVYDYIIPR | **ANDV-L4** | GLLENFMKGLPAAIP |
| **ANDV-N** | TEIATPHSVWVFACA | **ANDV-Gc** | RIVGPITAKVPSTSS | **ANDV-Gn** | PDGNTRDHVNLVLNR | **ANDV-L1** | LTVQGAYLPSTISSD | **ANDV-L2** | LKGMSGALQEDGHLA | **ANDV-L3** | HPVYDYIIPRNRDNL | **ANDV-L4** | FMKGLPAAIPPAAYS |
| **ANDV-N** | PHSVWVFACAPDRCP | **ANDV-Gc** | ITAKVPSTSSTDTLK | **ANDV-Gn** | RDHVNLVLNRDVSFQ | **ANDV-L1** | AYLPSTISSDQCNTR | **ANDV-L2** | GALQEDGHLAASVTL | **ANDV-L3** | YIIPRNRDNLLVYLV | **ANDV-L4** | PAAIPPAAYSLIMNK |
| **ANDV-N** | VFACAPDRCPPTALY | **ANDV-Gc** | PSTSSTDTLKGTAFA | **ANDV-Gn** | LVLNRDVSFQDLSDN | **ANDV-L1** | TISSDQCNTRIGCLE | **ANDV-L2** | DGHLAASVTLIEAIR | **ANDV-L3** | NRDNLLVYLVRKLND | **ANDV-L4** | PAAYSLIMNKAKISV |
| **ANDV-N** | PDRCPPTALYVAGVP | **ANDV-Gc** | TDTLKGTAFAGVPMY | **ANDV-Gn** | DVSFQDLSDNPCKVD | **ANDV-L1** | QCNTRIGCLEISKNQ | **ANDV-L2** | ASVTLIEAIRFLNRS | **ANDV-L3** | LVYLVRKLNDPSIIT | **ANDV-L4** | LIMNKAKISVDLFMF |
| **ANDV-N** | PTALYVAGVPELGAF | **ANDV-Gc** | GTAFAGVPMYSSLST | **ANDV-Gn** | DLSDNPCKVDLHTQA | **ANDV-L1** | IGCLEISKNQPETPV | **ANDV-L2** | IEAIRFLNRSQTNPN | **ANDV-L3** | RKLNDPSIITAMTMQ | **ANDV-L4** | AKISVDLFMFNELLA |
| **ANDV-N** | VAGVPELGAFFSILQ | **ANDV-Gc** | GVPMYSSLSTLVRNA | **ANDV-Gn** | PCKVDLHTQAIEGAW | **ANDV-L1** | ISKNQPETPVQMLAL | **ANDV-L2** | FLNRSQTNPNVIDMY | **ANDV-L3** | PSIITAMTMQSPLQL | **ANDV-L4** | DLFMFNELLALINRN |
| **ANDV-N** | ELGAFFSILQDMRNT | **ANDV-Gc** | SSLSTLVRNADPEFV | **ANDV-Gn** | LHTQAIEGAWGSGVG | **ANDV-L1** | PETPVQMLALDISYK | **ANDV-L2** | QTNPNVIDMYEQTKQ | **ANDV-L3** | AMTMQSPLQLRFRMQ | **ANDV-L4** | NELLALINRNNILNL |
| **ANDV-N** | FSILQDMRNTIMASK | **ANDV-Gc** | LVRNADPEFVFSPGI | **ANDV-Gn** | IEGAWGSGVGFTLTC | **ANDV-L1** | QMLALDISYKYLSLT | **ANDV-L2** | VIDMYEQTKQSKAQA | **ANDV-L3** | SPLQLRFRMQAKQHM | **ANDV-L4** | LINRNNILNLDGIEE |
| **ANDV-N** | DMRNTIMASKSVGTA | **ANDV-Gc** | DPEFVFSPGIIPESN | **ANDV-Gn** | GSGVGFTLTCTVGLT | **ANDV-L1** | DISYKYLSLTRDELI | **ANDV-L2** | EQTKQSKAQARIVRK | **ANDV-L3** | RFRMQAKQHMKVCRY | **ANDV-L4** | NILNLDGIEETSEGY |
| **ANDV-N** | IMASKSVGTAEEKLK | **ANDV-Gc** | FSPGIIPESNHSVCD | **ANDV-Gn** | FTLTCTVGLTECPSF | **ANDV-L1** | YLSLTRDELINYYSP | **ANDV-L2** | SKAQARIVRKYQRTE | **ANDV-L3** | AKQHMKVCRYEGEWV | **ANDV-L4** | DGIEETSEGYSTVTS |
| **ANDV-N** | SVGTAEEKLKKKSAF | **ANDV-Gc** | IPESNHSVCDKKTVP | **ANDV-Gn** | TVGLTECPSFMTSIK | **ANDV-L1** | RDELINYYSPRVHFQ | **ANDV-L2** | RIVRKYQRTEADRGF | **ANDV-L3** | KVCRYEGEWVTFREV | **ANDV-L4** | TSEGYSTVTSMSSKQ |
| **ANDV-N** | EEKLKKKSAFYQSYL | **ANDV-Gc** | HSVCDKKTVPITWTG | **ANDV-Gn** | ECPSFMTSIKACDLA | **ANDV-L1** | NYYSPRVHFQSSPNV | **ANDV-L2** | YQRTEADRGFFITTL | **ANDV-L3** | EGEWVTFREVLAAAD | **ANDV-L4** | STVTSMSSKQWSEEM |
| **ANDV-N** | KKSAFYQSYLRRTQS | **ANDV-Gc** | KKTVPITWTGYLPIS | **ANDV-Gn** | MTSIKACDLAMCYGS | **ANDV-L1** | RVHFQSSPNVKEPGT | **ANDV-L2** | ADRGFFITTLPTRVR | **ANDV-L3** | TFREVLAAADSFATS | **ANDV-L4** | MSSKQWSEEMSLMSD |
| **ANDV-N** | YQSYLRRTQSMGIQL | **ANDV-Gc** | ITWTGYLPISGEMEK | **ANDV-Gn** | ACDLAMCYGSTVANL | **ANDV-L1** | SSPNVKEPGTLKLGL | **ANDV-L2** | FITTLPTRVRLEIIE | **ANDV-L3** | LAAADSFATSYQPTE | **ANDV-L4** | WSEEMSLMSDDDIDD |
| **ANDV-N** | RRTQSMGIQLDQKII | **ANDV-Gc** | YLPISGEMEKVTGCT | **ANDV-Gn** | MCYGSTVANLARGSN | **ANDV-L1** | KEPGTLKLGLSQLNP | **ANDV-L2** | PTRVRLEIIEDYFDA | **ANDV-L3** | SFATSYQPTERDMDL | **ANDV-L4** | SLMSDDDIDDMEDFT |
| **ANDV-N** | MGIQLDQKIIILYML | **ANDV-Gc** | GEMEKVTGCTVFCTL | **ANDV-Gn** | TVANLARGSNTVKVV | **ANDV-L1** | LKLGLSQLNPLSKSI | **ANDV-L2** | LEIIEDYFDAIAKVV | **ANDV-L3** | YQPTERDMDLFNTLV | **ANDV-L4** | DDIDDMEDFTIALDD |
| **ANDV-N** | DQKIIILYMLSWGKE | **ANDV-Gc** | VTGCTVFCTLAGPGA | **ANDV-Gn** | ARGSNTVKVVGKGGH | **ANDV-L1** | SQLNPLSKSILDNVG | **ANDV-L2** | DYFDAIAKVVPEEYI | **ANDV-L3** | RDMDLFNTLVSCTFS | **ANDV-L4** | MEDFTIALDDIDFEQ |
| **ANDV-N** | ILYMLSWGKEAVNHF | **ANDV-Gc** | VFCTLAGPGASCEAY | **ANDV-Gn** | TVKVVGKGGHSGSSF | **ANDV-L1** | LSKSILDNVGKHKKD | **ANDV-L2** | IAKVVPEEYISYGGD | **ANDV-L3** | FNTLVSCTFSKEYAW | **ANDV-L4** | IALDDIDFEQINLEE |
| **ANDV-N** | SWGKEAVNHFHLGDD | **ANDV-Gc** | AGPGASCEAYSENGI | **ANDV-Gn** | GKGGHSGSSFKCCHD | **ANDV-L1** | LDNVGKHKKDKGLFG | **ANDV-L2** | PEEYISYGGDKKVLN | **ANDV-L3** | SCTFSKEYAWKDFLN | **ANDV-L4** | IDFEQINLEEDIQHF |
| **ANDV-N** | AVNHFHLGDDMDPEL | **ANDV-Gc** | SCEAYSENGIFNISS | **ANDV-Gn** | SGSSFKCCHDTDCST | **ANDV-L1** | KHKKDKGLFGEIIDS | **ANDV-L2** | SYGGDKKVLNIQNAL | **ANDV-L3** | KEYAWKDFLNEVRCE | **ANDV-L4** | INLEEDIQHFLQDES |
| **ANDV-N** | HLGDDMDPELRQLAQ | **ANDV-Gc** | SENGIFNISSPTCLV | **ANDV-Gn** | KCCHDTDCSTEGLLA | **ANDV-L1** | KGLFGEIIDSINVAS | **ANDV-L2** | KKVLNIQNALEKALR | **ANDV-L3** | KDFLNEVRCEVLTTR | **ANDV-L4** | DIQHFLQDESAYVGD |
| **ANDV-N** | MDPELRQLAQSLIDT | **ANDV-Gc** | FNISSPTCLVNKVQR | **ANDV-Gn** | TDCSTEGLLASAPHL | **ANDV-L1** | EIIDSINVASQIQIN | **ANDV-L2** | IQNALEKALRWASGV | **ANDV-L3** | EVRCEVLTTRHVHRP | **ANDV-L4** | LQDESAYVGDLLIQT |
| **ANDV-N** | RQLAQSLIDTKVKEI | **ANDV-Gc** | PTCLVNKVQRFRGSE | **ANDV-Gn** | EGLLASAPHLERVTG | **ANDV-L1** | INVASQIQINACAKI | **ANDV-L2** | EKALRWASGVSEITT | **ANDV-L3** | VLTTRHVHRPKIART | **ANDV-L4** | AYVGDLLIQTEDIEV |
| **ANDV-N** | SLIDTKVKEISNQEP | **ANDV-Gc** | NKVQRFRGSEQKINF | **ANDV-Gn** | SAPHLERVTGFNQID | **ANDV-L1** | QIQINACAKIIEQIL | **ANDV-L2** | WASGVSEITTSTGKS | **ANDV-L3** | HVHRPKIARTFTVRE | **ANDV-L4** | LLIQTEDIEVKKIRG |
| **ANDV-N** | DTKVKEISNQEPLKL | **ANDV-Gc** | FRGSEQKINFICQRV | **ANDV-Gn** | ERVTGFNQIDSDKVY | **ANDV-L1** | ACAKIIEQILSNLEI | **ANDV-L2** | SEITTSTGKSIKFKR | **ANDV-L3** | KIARTFTVREKDQAI | **ANDV-L4** | EDIEVKKIRGVTRVL |
| | | **ANDV-Gc** | QKINFICQRVDQDVV | **ANDV-Gn** | FNQIDSDKVYDDGAP | **ANDV-L1** | IEQILSNLEINIGEI | **ANDV-L2** | STGKSIKFKRKLMYV | **ANDV-L3** | FTVREKDQAIQNPIN | **ANDV-L4** | KKIRGVTRVLEPVKL |

| | | | | | | | | | | | | | |
|---|---|---|---|---|---|---|---|---|---|---|---|---|---|
| | | **ANDV-Gc** | ICQRVDQDVVVYCNG | **ANDV-Gn** | SDKVYDDGAPPCTFK | **ANDV-L1** | SNLEINIGEINASMP | **ANDV-L2** | IKFKRKLMYVSADAT | **ANDV-L3** | KDQAIQNPINSVIGY | **ANDV-L4** | VTRVLEPVKLLKSWV |
| | | **ANDV-Gc** | DQDVVVYCNGQKKVI | **ANDV-Gn** | DDGAPPCTFKCWFTK | **ANDV-L1** | NIGEINASMPSPNKT | **ANDV-L2** | KLMYVSADATKWSPG | **ANDV-L3** | QNPINSVIGYKYALT | **ANDV-L4** | EPVKLLKSWVSKGLA |
| | | **ANDV-Gc** | VYCNGQKKVILTKTL | **ANDV-Gn** | PCTFKCWFTKSGEWL | **ANDV-L1** | NASMPSPNKTTGVDD | **ANDV-L2** | SADATKWSPGDNSAK | **ANDV-L3** | SVIGYKYALTVDEVS | **ANDV-L4** | LKSWVSKGLAIDKVY |
| | | **ANDV-Gc** | QKKVILTKTLVIGQC | **ANDV-Gn** | CWFTKSGEWLLGILN | **ANDV-L1** | SPNKTTGVDDLLNKF | **ANDV-L2** | KWSPGDNSAKFRRFT | **ANDV-L3** | KYALTVDEVSDVLDS | **ANDV-L4** | SKGLAIDKVYNPIGI |
| | | **ANDV-Gc** | LTKTLVIGQCIYTFT | **ANDV-Gn** | SGEWLLGILNGNWIV | **ANDV-L1** | TGVDDLLNKFYDNEL | **ANDV-L2** | DNSAKFRRFTQAIYD | **ANDV-L3** | VDEVSDVLDSAFFPE | **ANDV-L4** | IDKVYNPIGIILMAR |
| | | **ANDV-Gc** | VIGQCIYTFTSLFSL | **ANDV-Gn** | LGILNGNWIVVVVLV | **ANDV-L1** | LLNKFYDNELGKYML | **ANDV-L2** | FRRFTQAIYDGLSDN | **ANDV-L3** | DVLDSAFFPESLSAD | **ANDV-L4** | NPIGIILMARYMSKT |
| | | **ANDV-Gc** | IYTFTSLFSLMPDVA | **ANDV-Gn** | GNWIVVVVLVVILIL | **ANDV-L1** | YDNELGKYMLSILRK | **ANDV-L2** | QAIYDGLSDNKLKCC | **ANDV-L3** | AFFPESLSADLQVMK | **ANDV-L4** | ILMARYMSKTYNFSS |
| | | **ANDV-Gc** | SLFSLMPDVAHSLAV | **ANDV-Gn** | VVVLVVILILSIIMF | **ANDV-L1** | GKYMLSILRKTAAWH | **ANDV-L2** | GLSDNKLKCCVVDAL | **ANDV-L3** | SLSADLQVMKDGVYR | **ANDV-L4** | YMSKTYNFSSTPLAL |
| | | **ANDV-Gc** | MPDVAHSLAVELCVP | **ANDV-Gn** | VILILSIIMFSILCP | **ANDV-L1** | SILRKTAAWHIGHLV | **ANDV-L2** | KLKCCVVDALRNIYE | **ANDV-L3** | LQVMKDGVYRELGLD | **ANDV-L4** | YNFSSTPLALLNPYD |
| | | **ANDV-Gc** | HSLAVELCVPGLHGW | **ANDV-Gn** | SIIMFSILCPRRGHK | **ANDV-L1** | TAAWHIGHLVRDITE | **ANDV-L2** | VVDALRNIYETEFFM | **ANDV-L3** | DGVYRELGLDISSPE | **ANDV-L4** | TPLALLNPYDLTELE |
| | | **ANDV-Gc** | ELCVPGLHGWATVML | **ANDV-Gn** | MFSILCPRRGHKKTV | **ANDV-L1** | IGHLVRDITESLIAH | **ANDV-L2** | RNIYETEFFMSRKLH | **ANDV-L3** | ELGLDISSPEVLKRI | **ANDV-L4** | LNPYDLTELESVVKG |
| | | **ANDV-Gc** | GLHGWATVMLLSTFC | | | **ANDV-L1** | RDITESLIAHAGLRR | **ANDV-L2** | TEFFMSRKLHRYIDS | **ANDV-L3** | ISSPEVLKRIAPLLY | **ANDV-L4** | LTELESVVKGWGETV |
| | | **ANDV-Gc** | ATVMLLSTFCFGWVL | | | **ANDV-L1** | SLIAHAGLRRSKYWS | **ANDV-L2** | SRKLHRYIDSMENHS | **ANDV-L3** | VLKRIAPLLYKAGRS | **ANDV-L4** | SVVKGWGETVNDRFK |
| | | **ANDV-Gc** | LSTFCFGWVLIPAVT | | | **ANDV-L1** | AGLRRSKYWSVHAYD | **ANDV-L2** | RYIDSMENHSDAVED | **ANDV-L3** | APLLYKAGRSRVVIV | **ANDV-L4** | WGETVNDRFKDLDIE |
| | | **ANDV-Gc** | FGWVLIPAVTLIILK | | | **ANDV-L1** | SKYWSVHAYDHGNVI | **ANDV-L2** | MENHSDAVEDFLAFF | **ANDV-L3** | KAGRSRVVIVEGNVE | **ANDV-L4** | NDRFKDLDIEAQTVV |
| | | **ANDV-Gc** | IPAVTLIILKCLRVL | | | **ANDV-L1** | VHAYDHGNVILFILP | **ANDV-L2** | DAVEDFLAFFSNGVS | **ANDV-L3** | RVVIVEGNVEGTAES | **ANDV-L4** | DLDIEAQTVVKEKGV |
| | | **ANDV-Gc** | LIILKCLRVLIFSCS | | | **ANDV-L1** | HGNVILFILPSKSLE | **ANDV-L2** | FLAFFSNGVSANVKG | **ANDV-L3** | EGNVEGTAESICSYW | **ANDV-L4** | AQTVVKEKGVQPEDV |
| | | **ANDV-Gc** | CLRVLIFSCSHYTNE | | | **ANDV-L1** | LFILPSKSLEVVGSY | **ANDV-L2** | SNGVSANVKGNWLQG | **ANDV-L3** | GTAESICSYWLKTMS | **ANDV-L4** | KEKGVQPEDVLPDSL |
| | | **ANDV-Gc** | IFSCSHYTNESKFKF | | | **ANDV-L1** | SKSLEVVGSYIRYFT | **ANDV-L2** | ANVKGNWLQGNLNKC | **ANDV-L3** | ICSYWLKTMSLIKTI | **ANDV-L4** | QPEDVLPDSLFSFRH |
| | | **ANDV-Gc** | HYTNESKFKFILEKV | | | **ANDV-L1** | VVGSYIRYFTVFKDG | **ANDV-L2** | NWLQGNLNKCSSLFG | **ANDV-L3** | LKTMSLIKTIRVRPK | **ANDV-L4** | LPDSLFSFRHVDVLL |
| | | **ANDV-Gc** | SKFKFILEKVKVEYQ | | | **ANDV-L1** | IRYFTVFKDGIGLID | **ANDV-L2** | NLNKCSSLFGAAVSL | **ANDV-L3** | LIKTIRVRPKKEVLK | **ANDV-L4** | FSFRHVDVLLRRLFP |
| | | **ANDV-Gc** | ILEKVKVEYQKTMGS | | | **ANDV-L1** | VFKDGIGLIDADNID | **ANDV-L2** | SSLFGAAVSLLFREV | **ANDV-L3** | RVRPKKEVLKAMSLY | **ANDV-L4** | VDVLLRRLFPRDPVS |
| | | **ANDV-Gc** | KVEYQKTMGSMVCDV | | | | | | | | | **ANDV-L4** | LLRRLFPRDPVSTFY |
| | | **ANDV-Gc** | KTMGSMVCDVCHHEC | | | | | | | | | | |
| | | **ANDV-Gc** | MVCDVCHHECETAKE | | | | | | | | | | |
| | | **ANDV-Gc** | CHHECETAKELESHR | | | | | | | | | | |
| | | **ANDV-Gc** | ETAKELESHRQSCIN | | | | | | | | | | |
| | | **ANDV-Gc** | LESHRQSCINGQCPY | | | | | | | | | | |
| | | **ANDV-Gc** | QSCINGQCPYCMTIT | | | | | | | | | | |

| | | | |
|---|---|---|---|
| | | **ANDV-Gc** | GQCPYCMTITEATES |
| | | **ANDV-Gc** | CMTITEATESALQAH |
| | | **ANDV-Gc** | EATESALQAHYSICK |
| | | **ANDV-Gc** | ALQAHYSICKLTGRF |
| | | **ANDV-Gc** | YSICKLTGRFQEALK |
| | | **ANDV-Gc** | LTGRFQEALKKSLKK |
| | | **ANDV-Gc** | QEALKKSLKKPEVKK |
| | | **ANDV-Gc** | KSLKKPEVKKGCYRT |
| | | **ANDV-Gc** | PEVKKGCYRTLGVFR |
| | | **ANDV-Gc** | GCYRTLGVFRYKSRC |
| | | **ANDV-Gc** | LGVFRYKSRCYVGLV |
| | | **ANDV-Gc** | YKSRCYVGLVWCLLL |
| | | **ANDV-Gc** | YVGLVWCLLLTCEIV |
| | | **ANDV-Gc** | WCLLLTCEIVIWAAS |
| | | **ANDV-Gc** | TCEIVIWAASAETPL |
| | | **ANDV-Gc** | IWAASAETPLMESGW |
